# High-plex Multiomic Analysis in FFPE at Subcellular Level by Spatial Molecular Imaging

**DOI:** 10.1101/2021.11.03.467020

**Authors:** Shanshan He, Ruchir Bhatt, Carl Brown, Emily A. Brown, Derek L. Buhr, Kan Chantranuvatana, Patrick Danaher, Dwayne Dunaway, Ryan G. Garrison, Gary Geiss, Mark T. Gregory, Margaret L. Hoang, Rustem Khafizov, Emily E. Killingbeck, Dae Kim, Tae Kyung Kim, Youngmi Kim, Andrew Klock, Mithra Korukonda, Alecksandr Kutchma, Zachary R. Lewis, Yan Liang, Jeffrey S. Nelson, Giang T. Ong, Evan P. Perillo, Joseph C. Phan, Tien Phan-Everson, Erin Piazza, Tushar Rane, Zachary Reitz, Michael Rhodes, Alyssa Rosenbloom, David Ross, Hiromi Sato, Aster W. Wardhani, Corey A. Williams-Wietzikoski, Lidan Wu, Joseph M. Beechem

## Abstract

The Spatial Molecular Imaging platform (CosMx^TM^ SMI, NanoString Technologies, Seattle, WA) utilizes high-plex *in-situ* imaging chemistry for both RNA and protein detection. This automated instrument provides 1000’s of plex, at high sensitivity (1 to 2 copies/cell), very low error rate (0.0092 false calls/cell) and background (∼0.04 counts/cell). The imaging system generates three-dimensional super-resolution localization of analytes at ∼2 million cells per sample, four samples per run. Cell segmentation is morphology-based using antibodies, compatible with FFPE samples. Multiomic data (980 RNAs, 108 proteins) were measured at subcellular resolution using FFPE tissues (non-small cell lung (NSCLC) and breast cancer) and allowed identification of over 18 distinct cell types, 10 unique tumor microenvironments, and 100 pairwise ligand-receptor interactions. Over 800,000 single cells and ∼260 million transcripts data are released into the public domain allowing extended data analysis by the entire spatial biology research community.

## Introduction

Understanding the spatial distribution of RNA and protein in tissues has a great potential to expand our knowledge of all aspects of life science research (1–6). Fluorescence *in-situ* hybridization (FISH) and immunohistochemistry (IHC) are the most traditional technologies to assess the spatial distribution of RNA and protein in fixed tissue samples (7–10). However, these technologies can only detect a handful of targets at a time. In addition, high-plex, high sensitivity multiomic detection (RNA and proteins simultaneously) in formalin-fixed, paraffin-embedded (FFPE) has not yet been accomplished. Current multiplex IHC techniques (11) involve a long assay time, which is not easily automated, and often require extensive assay optimization, hindering their establishment as routine methods.

For high-throughput high-plex RNA profiling with single-cell resolution, the most commonly used technology to date is single-cell RNA-seq (scRNA-seq), such as Drop-seq, inDrop, and Chromium^TM^ (10X Genomics) (12–15). These technologies require tissue dissociation and cell isolation into nanoliter droplets with hydrogel beads bearing barcoding DNA primers. Although these methods enable whole transcriptome RNA profiling at single-cell resolution, the lack of spatial information limits this approach to generating a very detailed “parts-list” of the tissue.

Recently, multiple spatially resolved technologies have been developed that attempt to maximize the number of markers observable at the same time. These methods can be classified into two categories: 1) Profilers that enable spatially resolved high-plex data, often at the whole transcriptome level, from small sub-regions of the tissue using next-generation sequencing (NGS) as a readout. Profilers are not single-cell resolved and often have RNA capture reagents arranged in a pre-defined grid-like pattern on which tissue sections are mounted and 2) Imagers that provide true single-cell and subcellular resolution using *in-situ* reagents, though a plex level is smaller than the whole transcriptome in most platforms.

Profiling-based techniques include Spatial Transcriptomics (16), Slide-seqV2 (17), pixel-seq (18), dbit-seq (19), and Digital Spatial Profiling (20) that offer the ability to perform highly multiplexed profiling based on the detection readout using NGS. These techniques are limited, however, by not being resolved at the single-cell level and often suffer from dead-space areas in the sample where no measurements are performed. Although the active RNA capture regions (“spots”) can be precisely organized, the placement of the tissue sample onto the spots has limited control in the selection of regions to be analyzed, ignoring the morphological information in the tissue. Also, the spot size in the RNA capture area introduces analysis issues; for example, if the spot is set to a larger size (50 μm), many cells are randomly selected by the spot location on tissue. If the spots are small (≤ 2 μm), the number of captured transcripts is very low and the spots are grouped together for analysis, leading to the same issues as larger spots. In addition, many RNA profiling methods capture RNA molecules via the poly-A tail, which is inefficient for the subsequent sequencing process since poly-A transcripts are dominated by highly expressed genes (21). As spot-based methods rely on the mobility of RNA molecules in the tissue sample, they struggle to work with FFPE samples without requiring separate chemistries for different sample types. The *in-situ* solution offered by the GeoMx® Digital Spatial Profiler (DSP) overcomes many of these issues, providing a method for highly multiplexed spatial profiling of RNA and proteins on FFPE and frozen samples (20).

Single-cell high-plex imagers use sequential cycles of probe hybridization and imaging and offer the potential to combine the benefits of scRNA-seq analysis with added spatial resolution at single-cell or even subcellular resolution. Recent single-cell high-plex imaging technologies include MERFISH (22, 23), Molecular Cartography™ (24), FISSEQ (25), and seqFISH+ (26). Despite isolated proof-of-concept experiments demonstrating RNA-plex levels in ∼10,000 targets (26, 27), most molecular imaging systems allow lower plex from 100 (24) to 500 (22) targets. As these methods were developed for fresh-frozen samples, the detection efficiency can be very low on FFPE tissue samples.

For protein detection, CODEX and InSituPlex® methodologies increase target-plex by labeling antibodies with unique oligonucleotide tags that are identified with fluorescence detection during amplification of the probe sequence or through cyclical hybridization of fluorescently labeled reverse-complement sequences (28). Although cyclic IF and ion-beam imaging have enabled high-plex detection of dozens of proteins *in-situ* (29, 30), both approaches are limited in the maximal plex number due to on-instrument time or availability of metal isotopes.

Here, we describe the chemistry and applications of a Spatial Molecular Imager (SMI) to address many of the current unmet needs in spatial high-plex profiling of RNA and protein expression. SMI is a completely automated and integrated platform comprising chemistry, hardware, and software that enables highly sensitive spatial profiling of 980 RNAs and 108 proteins in FFPE tissues at single-cell and subcellular resolution.

### 1. Spatial Molecular Imager (SMI) Chemistry and Workflow

SMI is developed to perform multiple cycles of the nucleic-acid hybridization of fluorescent molecular barcodes to enable *in-situ* measurement of RNAs and proteins on intact biological samples at subcellular resolution. The SMI chemistry is an enzyme-free, nucleic acid amplification-free, hybridization-based single-molecule barcode sequencing methodology, which can be directly applied to intact FFPE and fresh-frozen tissues on 1 mm-thick standard glass slides for pathology.

The SMI chemistry relies on *in-situ* hybridization (ISH) probes and fluorescent readout probes called reporters (Figure 1A) to detect RNAs in intact tissue. The ISH probes consist of a DNA of length 35 to 50 nucleotides (nt) that will hybridize with the RNA target in the tissue (target-binding domain), coupled with a stretch of 60-80 nt of DNA “readout domain” for RNA identification. The readout domain consists of four consecutive 10-20 nt sequences that allow four individual SMI imaging barcodes (reporters) to bind sequentially. In addition, for each gene, five RNA-detection oligonucleotide probes (“tiles”) of ISH probes were designed to detect different regions of the RNA target, and each tile can independently record RNA-location identity.

**Figure 1.**
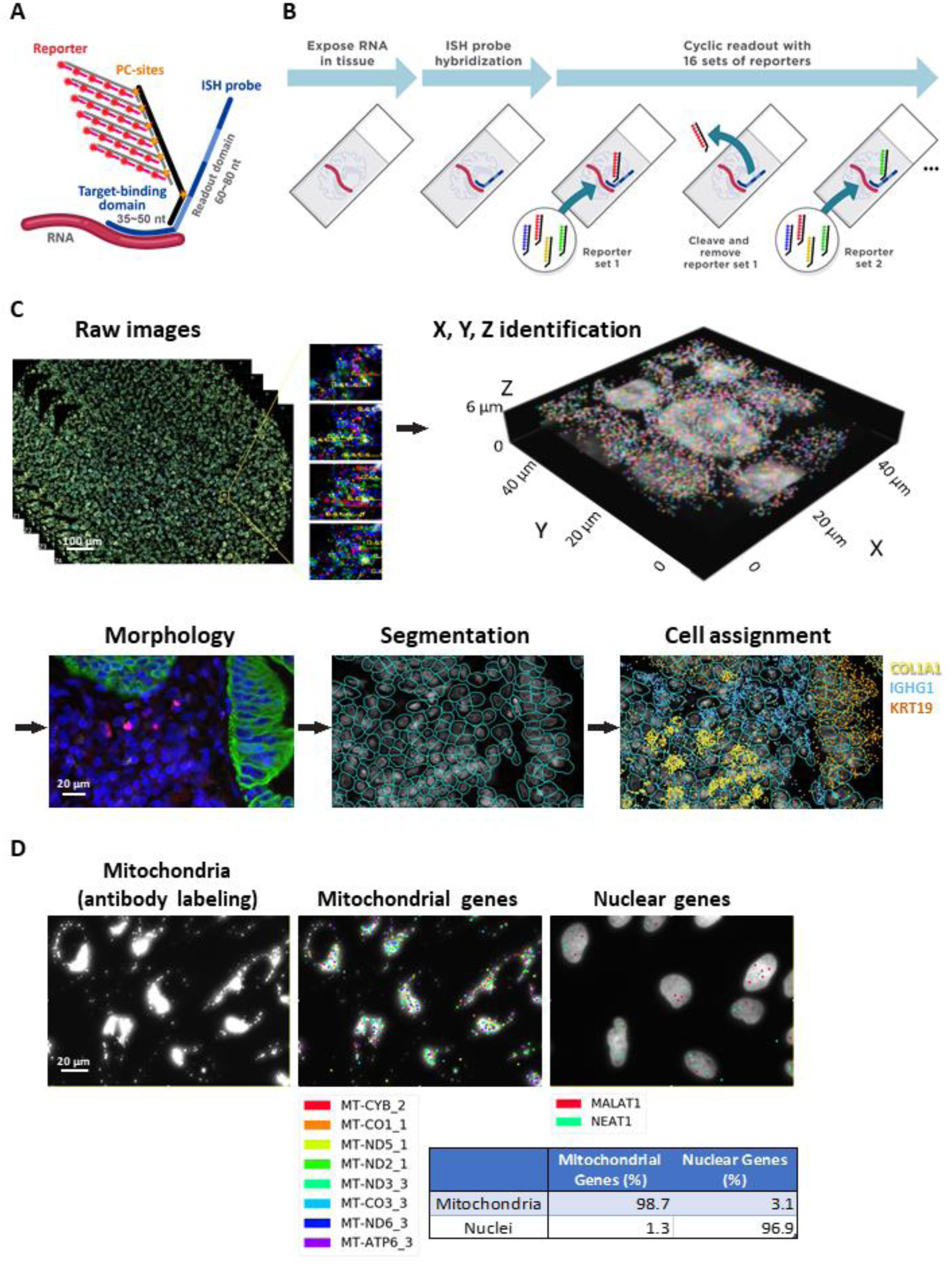
CosMx SMI Chemistry and Workflow and 3D Mapping of RNA at Single-Cell Subcellular Resolution Using Morphology-Based Cell Segmentation. **A.** Schematic description of the SMI ISH probe and reporter design. The ISH probe consists of the target-binding domain and readout domain. The target-binding domain is a 35-50 nt DNA sequence that hybridizes with target RNA. The readout domain is a 60-80 nt DNA sequence that contains 4 consecutive 10-20 nt reporter-landing domains, where each landing domain can be hybridized with a unique reporter. With a 64-bit barcoding design, there are a total of 64 unique reporter-landing sequences. Each reporter is a single-color branched structure assembled with oligos that are conjugated with one of the four fluorophores and will be detected as one of four colors (blue, green, yellow, or red) in SMI images. Each reporter has a controlled number of 15-60 dyes with 6 photocleavable (PC) sites to efficiently quench signals by UV illumination and a washing step prior to each cyclic reporter readout. **B.** Illustration of the SMI RNA assay workflow. The FFPE slide undergoes standard tissue preparation to expose RNA targets for ISH probe hybridization. The sample is assembled into a flow cell and loaded onto an SMI instrument for cyclic readout with 16 sets of reporters. As each reporter set contains 4 reporters with 4 different fluorophores, a total of 64 unique reporters are used in the SMI assay to bind to the different reporter landing domains on ISH probes. Following each set of reporter hybridization, high-resolution Z-stacked images are acquired, followed by cleavage and removal of fluorophores from the reporters prior to incubation with the next set of reporters. C. Analysis workflow to transform raw images to decoded RNA transcripts at subcellular resolution. The workflow includes: 1) three-dimensional primary image processing to identify and register reporter spots, 2) decoding of reporter spots to RNA transcripts with registered X, Y, Z spatial location, 3) outlining of nuclei and cell boundaries with DAPI and antibodies after cyclic reporter readout for morphology-based cell segmentation, and 4) assigning RNA transcripts to single cells. As an example of the final output of the analysis workflow, three identified genes (COL1A1 [yellow], IGHG1 [cyan], and KRT19 [orange]) in FFPE human lung tissue are overlayed with the segmented cells based on their registered spatial information. Scale bars: 100 μm for raw images, 20 μm for the images with morphology-based cell segmentation images. D. Demonstration of subcellular resolution in U2OS cells. Transcripts included in a 980-plex panel located in mitochondria and nuclei are shown. These genes were read out on fixed U2OS cells with the SMI RNA assay, followed by labeling with morphology antibodies for membrane (CD298) and mitochondria, as well as DAPI stain for nuclei. Scale bar: 20 μm. The table shows the quantification of transcripts detected in these two cell compartments.

Each tile of the ISH probe has its unique sequence in the target-binding domain whereas all tiles share the same sequence in the readout domain. This design enables the highest detection sensitivity in FFPE tissue, in which RNA targets may be highly fragmented. For instance, even if only one of the tiles successfully binds to the target, the RNA target can be detected at the readout step. Each reporter construct contains the controlled number of 15-to-60 dyes, depending on desired sensitivity, assembled with fluorophore-conjugated oligos with photocleavable (PC)-linkers (Figure 1A). All reporters are single-color, containing one of the four fluorophores: Alexa Fluor-488, ATTO 532, Dyomics-605, or Alexa Fluor-647. The key advantages of the SMI reporter chemistry are high signal-to-noise ratio (SNR) detection over the background for accurate spot calling and fast fluorescent signal quenching by ultraviolet (UV) cleavage of PC-linkers (Figure 1B).

For sample preparation, SMI utilizes standard methods performed for FISH on FFPE tissue sections to expose RNA targets, followed by the introduction of fluorescent bead-based fiducials that are fixed to the tissue to provide an optical reference for cyclic image acquisition and registration. Following hybridization of ISH probes, slides are washed, assembled into a flow cell, and placed within a fluidic manifold on the SMI instrument for RNA readout and morphological imaging. In RNA assay readout, the tissue is hybridized with 16 sets of fluorescent reporters sequentially; each reporter set contains four single-color reporter pools. The reporters specifically bind to ISH probes during the 16 rounds of reporter hybridization according to the barcode assigned to each gene (Figure S1). After the incubation of each set of reporters, high-resolution Z-stacked images are acquired for downstream analysis (Figure 1C). Prior to the incubation with the next set of reporters, PC-linkers are cleaved by UV illumination and released fluorophores are removed by washing (Figure 1B).

The SMI encoding scheme is designed to assign a unique barcode to each target transcript from a set of 64-bit barcodes (4 color reporters in each readout round over 16 readout rounds) with the Hamming distance 4 (HD4) and Hamming weight 4 (HW4) (31). Every barcode is separated by an HD of at least 4 from all other barcodes to maximally suppress RNA decoding error. Every barcode has a constant HW4, in which each target is “on” in 4 rounds and “off” in 12 rounds during the 16 rounds of reporter hybridization. This “on” and “off” signal barcode design allows for continued expansion to even higher plex since only a subset of RNAs is “on” in any given cycle. Note that the nature of these on-off imaging signals represents a “deterministic super-resolution imaging system”, and hence each SMI imaging barcode can be located well below the diffraction limit of the imaging system. For each reporter hybridization round, a single reporter can bind to one of the reporter-landing domains on the ISH probe of the gene (Figure 1A and B, Figure S1). The 64-bit encoding scheme with the HD4 and HW4 yields 1,210 barcodes, from which a subset of 980 barcodes is selected to detect 960 target genes and 20 negative control targets (Table S1). Negative control probes are modeled after synthetic sequences from the External RNA Controls Consortium (ERCC) (22). Since they target alien sequences that are non-existent in human tissue, negative control probes serve as non-target controls for quantification of non-specific ISH probe hybridization. Five RNA-detection oligonucleotide probes/tiles are designed per gene and negative control target (Table S2). The colors of these barcodes are randomly assigned to targets to avoid any color-code-induced bias (Table S3). The remaining 230 barcodes were left as blank controls for misidentification quantification of reporter readout.

Another key component of the SMI platform is the integrated and fully automated fluidic and imaging capability. SMI features a large scan area (range 16 mm^2^ to 375 mm^2^) on each tissue slide and supports up to 4 slides simultaneously. The run-time per sample is dependent on the number of flow cells utilized, the area imaged per sample, and the size of the target assay panel. The on-instrument time for the commercial-release SMI instrument will range from four samples per day for ∼16 mm^2^ tissue area to ∼0.5 per day for 100 mm^2^ samples. The ability to measure four slides per single SMI run, with each slide allowing multiple samples to be placed onto a single slide (375 mm^2^ active imaging area), provides extreme flexibility in the maximized throughput of the SMI system.

The RNA and protein SMI imaging barcodes over-background pixels are located (in 1 of the 4 color channels), and the intensity distribution of each barcode is fit using a 2D parabola at sub-pixel resolution. Images from different cycles and different colors are registered using an affine transform to align the fluorescent bead reagents added to each tissue slide. The positions of the imaging spots contribute to an individual RNA target call by generating a very rich super-resolution data matrix of the final X, Y, and Z coordinates of the analyte. The position of each transcript call made using SMI can be estimated, on average, within a disc of radius ∼50 nm by making use of the super-resolution locations of individual reporter binding events contributing to that transcript call over the timescale for the entire experiment (Figure S2). Z-axis localization is dependent on the number of optical Z-stacks acquired during data collection. The step sizes of 800 nm were used for data collection; therefore, Z-axis localization is approximately +/- 400 nm. Unlike XY axes, there was no fitting in the Z-axis with different planes, and accuracy is close to pixel-size or step-size resolution.

### 2. 3D RNA Mapping and Cell Segmentation at Subcellular Resolution

The combination of the SMI chemistry, hardware, and software enable high-plex spatial RNA profiling within single cells with very high sensitivity. For spatial profiling, the data of fluorescent signals in Z-stacked raw images are transformed into decoded RNA transcript information at their registered spatial location in three dimensions (3D) and assigned to single cells that are identified by segmentation (Figure 1C). During the primary image processing, the fluorescence signals in raw images are transformed into digital output comprising of detected “spots” localized in X-, Y-, and Z-dimensions for subsequent RNA decoding. A spot is identified in an image as an isolated fluorescence signal with the intensity much higher than its neighbors. In biological tissues, however, dense fluorescent spots can be detected at a small spatial region, which may limit the accuracy of resolving each spot. To solve this potential issue, we developed custom image analysis algorithms to process 3D multi-channel image stacks obtained in each field of view (FOV). The key objective of this analytical method is to reduce the multi-dimensional image stack to a single list of individual reporter-binding events. This process is performed across all FOVs, concurrently with image acquisition during cyclic reporter readout. All spots pertaining to a given FOV are collated into a single list of XYZ locations of all individual reporter-binding events. This list is used in the next step to decode the gene-specific barcodes formed by these reporter-binding events.

The decoding algorithm enables the detection of as many transcripts as possible in crowded reporter-binding events while limiting the rate of error calls that can be generated due to the presence of multiple transcripts in close vicinity. In this algorithm, each unique XYZ location with at least one reporter-binding event is considered a ‘seed’ and used to construct a neighborhood in which gene-specific barcodes are searched. Two passes through of the data with the seed-centered neighborhood search are used to obtain a list of potential transcripts with their spatial locations. In the first pass-through of the data, the neighborhood search is limited to a radius of 0.5 pixels (90 nm). Because every gene barcode has 4 “on” spots, any seeds with fewer than 4 unique reporter probe-binding events in the neighborhood are removed from consideration for transcript decoding due to their inability to form a complete gene-specific barcode. All possible four reporter combinations of unique reporter probes in a seed’s neighborhood are then matched against a table of all potential gene-specific barcodes to detect the presence of a gene in a seed’s neighborhood. If more than one gene was detected in a seed’s neighborhood, the seed and all the transcripts detected in its neighborhood are dropped from further analysis. The impact of filtering out seeds with multiple target calls was minor (3-10%) on overall transcript calls.

Any reporter-binding events used in making transcript calls were removed from the dataset, and the process is repeated with a slightly increased search radius of 0.75 pixels to try to recover any potential transcript calls that may have been lost by local tissue motion. The transcript calls generated by these two passes through the data were further filtered to remove any potential duplicate calls or calls in neighborhoods with a high probability of making a transcript call by random chance. These various filtering steps are crucial to address potentially elevated error-call rates and duplicate calls of individual transcripts, ensuring that at the completion of the transcript decoding process, we only retain high-confidence transcript calls while maximizing the number of decoded transcripts.

Following RNA decoding and 3D spatial registration of target signals, the cell segmentation process is performed on tissue sections of the FFPE human lung to define cell boundaries based on tissue morphology markers (Figure 1C).

In this cell segmentation process, tissue is stained with a nuclear dye (DAPI) and labeled with antibodies for morphology markers, such as membrane (CD298), epithelial cells (pan-cytokeratin [PanCK]), and T cells (CD3). Defining accurate cell boundaries by segmentation is critical to data quality because it affects the spatial assignment of transcripts to specific cells. It is challenging to achieve high accuracy on tissue sections in which cells are tightly packed, shared 3D boundaries, or unevenly stained with morphology markers. To address this issue, we developed a cell segmentation pipeline that combines image preprocessing with machine learning for better accuracy (Figure S3 and S4). The signals of membrane channels were combined and normalized to the range of the nuclear channel, and then, image subtraction was performed between the nuclear channel and the normalized membrane image. After these preprocessing steps, the images were fed into pre-trained Cellpose neural network models (18). The Cellpose model has been validated with an average precision metric of 0.91 (*i.e.,* a 9% error) at a threshold level of 0.5 on the specialized data. In this study, we augmented the pure protein-based (cytoplasmic) Cellpose algorithm with an additional independent analysis of SMI nuclear-based signal using the Cellpose algorithm for a more accurate final cell-segmentation. Using this improved approach, we estimate the net segmentation error rate to be approximately 5% to 9%.

In the registered image, each transcript location is mapped to the corresponding cell and within the cellular compartment (*e.g.,* nuclei, cytoplasm, membrane). To demonstrate accurate RNA detection at the subcellular resolution, we designed a set of SMI probes that target mitochondrial genes and nuclear genes (32, 33). The cell compartments in U2OS cells were labeled with fluorescent probe-conjugated antibodies for mitochondria and membrane (CD298) as well as stained with DAPI for nuclei (Figure 1D). RNA profiling data shows that 98.7% of detected mitochondrial genes specifically colocalized with the antibody-based mitochondrial segment and 96.9% of nuclear genes strictly located within nuclei. These results indicate high accuracy in RNA detection and its spatial assignment at subcellular resolution.

### 3. SMI 980-plex RNA Panel and Cross-Platform Validation

For RNA profiling, we designed a panel targeting 980 genes to investigate the biology of single cells or subcellular compartments. Among those genes, 211 genes were selected for use in cell type identification, and the remaining genes were designed to capture critical cell states, cell-cell interactions, and hormone activity (Figure S5, list of genes in Table S1). The panel also includes 20 ERCC negative targets containing hybridization regions that are not complementary to the human genome or transcriptome (34). The panel design and gene selection process are fully described in the Methods.

To ensure that this panel was capable of quantifying RNAs in a wide range of biological samples, we profiled the SMI 980-plex panel on 16 different cell lines (CCRF-CEM, COLO205, DU145, EKVX, HCT116, HL60, HOP92, HS578T, IGROV1, M14, MDA-MB-468, MOLT4, PC3, RPMI-8226, SKMEL2, SUDHL4), spanning a broad range of biology and expression patterns (Table S4). Furthermore, in order to understand how SMI performs on “real-world” FFPE samples, the 980-plex assay was run on eight FFPE slides from five non-small cell lung cancer (NSCLC) tumors (Table S5). All samples were from tumor tissues archived in 2017 and 2018, the typical age for many archived cancer samples available to researchers (Table S6). The RNA in these samples ranged from “too degraded to sequence” to “medium quality to sequence” according to either RIN index or DV200 scoring (Table S7, Figure S6). These data, therefore, represent a range of sample types, including the most difficult class of archived material any researcher is likely to examine.

Using the SMI image processing methodology, hundreds of transcripts per cell were detected in the tested FFPE cell-line pellets with a maximum of 4,500 transcripts resolved in a single cell (Figure 2A, Table S4). Testing in FFPE NSCLC tissues identified an average of 260 counts per cell and a maximum of 2,737 transcripts (Figure 2B, Table S5). SMI RNA profiling also features a cell drop-out rate (cells contain fewer than 20 total transcripts) of less than 3% in the 16 cell lines tested. Background signals for the 20 ERCC control targets were extremely low (∼0.04 counts per target per cell), resulting in 12.3% of transcripts called being off-target background. Our single-cell distribution analysis revealed that even low expressing targets were detected well above the background (Figure 2A and B), confirming the high sensitivity of the SMI RNA assay.

**Figure 2.**
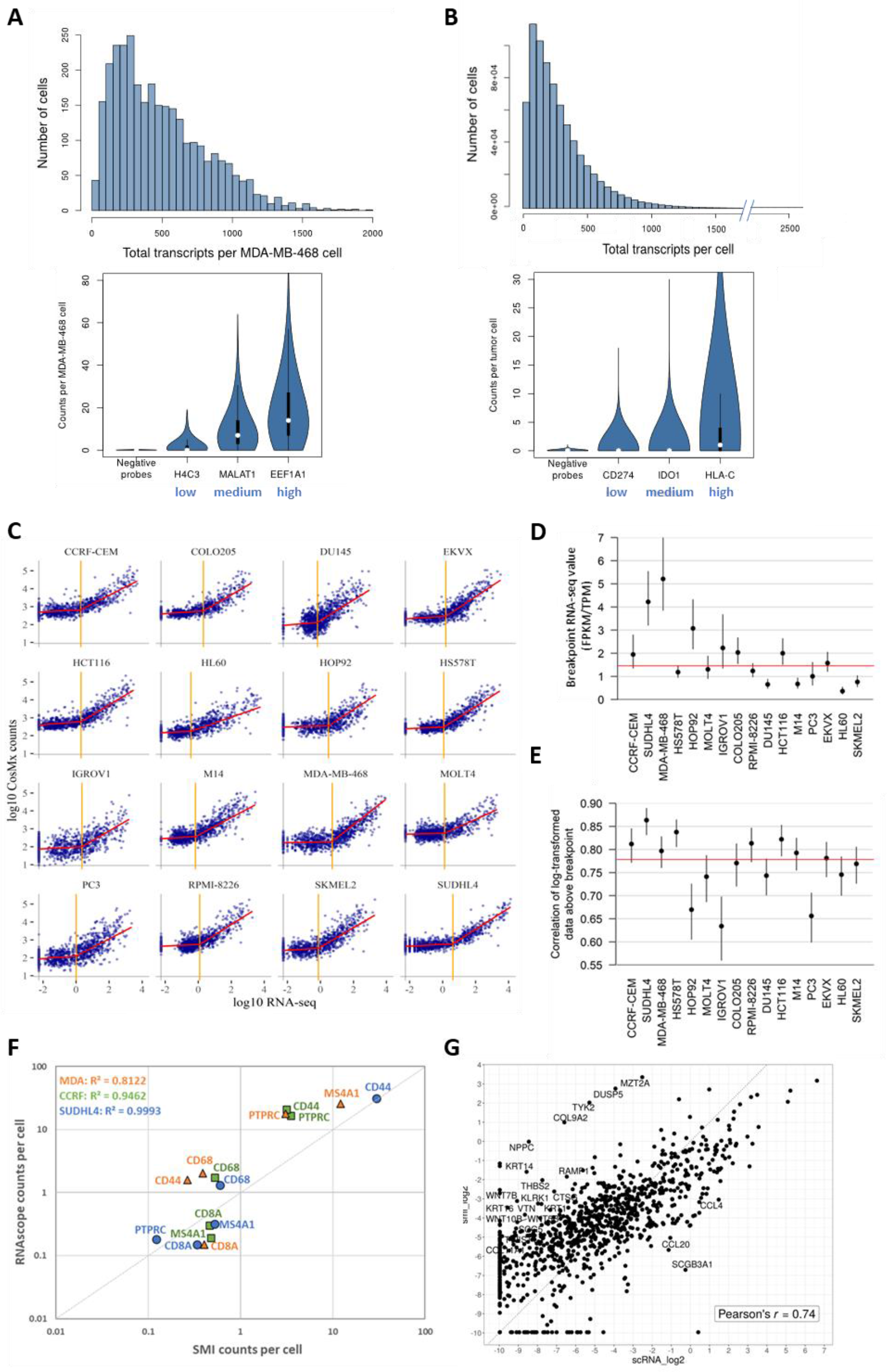
Single-Cell Distribution and Detection Sensitivity of Low, Medium, and High Expressers and Comparison of SMI Data to RNA-seq and RNAscope Data. **A.** Single-cell distribution of total transcripts per cell in FFPE MDA-MB-468 cell line is shown. Average counts per cell of genes representing low (H4C3), medium (MALAT1), high (EEF-1A1) expressers in this cell line and negative probes. **B.** Single-cell distribution of total transcripts per cell in FFPE lung tissue is shown. Among only tumor cells, average counts per cell of genes representing low (CD274), medium (IDO1), high (HLA-C) expressers and negative targets. **C.** For 16 cell lines, RNA-seq values are plotted against raw SMI counts. RNA-seq values are in TPM for cell lines from the NCI-60, and in FPKM for cell lines from the CCLE. Orange lines show breakpoints from segmented regression. Red lines show segmented regression fit. **D.** Estimated breakpoints and 95% confidence intervals from each cell line in FPKM or TPM depending on cell line origin. The red line shows the median value. **E.** Estimates and 95% confidence intervals for correlation between log-transformed RNA-seq and SMI data of genes with RNA-seq values above the breakpoint. The red line shows the median value. **F.** The average counts per cell of five genes (CD8A, CD44, CD68, RTPRC, MS4A1) are compared between RNAscope data and SMI data from three representative cell lines (MDA, SUDHL4, CCRF). **G.** For the SMI NSCLC dataset, the log2 of mean counts per gene in immune cell populations (100,193 cells) are compared to 215,423 cells in a published scRNA-seq dataset in NSCLC (40).

For cross-platform validation, SMI RNA counts were compared to bulk RNA-seq FPKM values (fragments per kilobase of transcript per million mapped reads) published in the Cancer Cell Line Encyclopedia (CCLE) (35) and TPM (transcripts per million) published as part of the NCI-60 Human Tumor Cell Lines Screen (36, 37). Segmented regression was fit predicting raw SMI counts from RNA-seq (Figure 2C). For the average cell line, the breakpoint in the segmented regression occurred at an RNA-seq value ranging from 0.36 to 5.2 (median: 1.443) (Figure 2D). Above this breakpoint, SMI counts linearly tracked RNA-seq counts with high correlation (Figure 2C). The results show adequate concordance between SMI counts and bulk RNA-seq values (Figure 2E), indicating that the panel can measure a wide range of biological processes. Furthermore, the comparison of SMI data from three representative cell lines to that of a conventional hybridization-based assay, RNAscope, revealed that the average counts of five representative genes per cell were highly correlated between these platforms (R^2^ = 0.8122 to 0.9993, Figure 2F). SMI counts per cell were also compared to the published CPM-normalized scRNA-seq counts (38) from three matched cancer cell lines (Pearson’s *r* = 0.74 to 0.76, Figure S7A). In addition, SMI data was compared to this scRNA-seq dataset for the fraction of cells that have non-zero counts for each of the 788 genes that are common in both datasets (Figure S7B). This fraction for each gene indicates the sensitivity ratio between the two technologies, as recently described in (39). The data points above the slope=1 line in a scatter plot would indicate that one technology has higher sensitivity than the other. In the three cell lines shown, the ratios of sensitivities of SMI over scRNA-seq are 520/268 in the EKVX lung adenocarcinoma cell line, 609/179 in the SKMEL2 melanoma cell line, and 569/219 in the T47D breast cancer cell line.

For the NSCLC dataset, SMI counts were compared to a published scRNA-seq dataset of CD45^+^ cells in NSCLC (40). The mean counts for the 952 shared genes in the immune cell population (100,193 cells) of the SMI dataset were compared to 215,423 cells in the scRNA-seq dataset. In the SMI data, the mean of the negative targets was subtracted from each gene’s mean counts. All counts from SMI and scRNA-seq were thresholded up to 1e-3 to enable log transformation and ensure positive numbers after subtracting the background from the SMI counts. The counts were log2 transformed and all 952 genes are plotted in Figure 2G (Pearson’s *r* = 0.74). For the average gene, the log2 ratio of the SMI signal strength over scRNA-seq signal strength is 0.61; that is, background-subtracted SMI data shows a 2^0.6^ = 1.51-fold stronger signal than scRNA-seq. For the genes with counts in the 25^th^ percentile and 75^th^ percentile for scRNA-seq, the average log2 ratio is 2.62 and -1.42, respectively. In addition, we compared the SMI bulk profile of each NSCLC sample to the log_10_ mean FPKM of NSCLC data in TCGA using the breakpoint regression used in Figure 2C. The breakpoints of five tissues, above which signal predominates over a background, occurred between 3.3 and 9.6 FPKM (Figure S7C). Correlations above the breakpoints ranged from 0.57 to 0.75. A small number of high outlier genes are evident, such as MALAT1, MZT2A, COL9A2, NPPC, DUSP5, and TYK2. These outliers were all a log_10_-unit above the trend line in 3 to 4 of the 5 samples. The outliers could be explained by sample variation due to different cell type composition and tumor states used for the public RNA-seq dataset. The recurring outliers can be partially due to platform effects such as different target accessibility for detection, or due to different sample preparation procedures for the *in-situ* imaging-based platform versus bulk RNA-seq. Most outliers in Lung 6 are keratin-encoding genes and only appear in this sample but not in others. This indicates that SMI procedures have only slight biases in gene quantification.

### 4. RNA-Based Cell Typing and Interaction Analysis in NSCLC

Measurements of eight slides from these NSCLC samples represent a total of 800,327 cells, of which nearly 96% of cells passed quality control, yielding 769,114 analyzable cells (Table S5). An average of 260 transcripts per cell were detected (Table S5 and S8), and 850 of the 960 potential genes in this panel were measured above the background (Table S5). A first step in the analysis of these datasets is to identify the high-level cell types of as many spatially resolved cells as possible. Single-cell expression profiles were derived by counting the transcripts of each gene that fell within the area assigned to a cell by the cell segmentation algorithm (Figures S3 and S4). An average of 4% of cells contained fewer than 20 total transcripts (Table S5), which were omitted from further analysis as this class of cells behaves anomalously when projected into lower dimensions (*e.g.*, UMAP). The average falsecode had 0.0092 counts in the average cell, indicated as the mean false calls/cell/target in Table S5.

Cell type in NSCLC tissue was determined by comparing individual cells’ expression profiles to reference profiles for different cell types (scRNA-seq and bulk RNA-seq of flow-sorted blood and stroma databases) (41), assigning each cell to the cell type under which its profile was most likely (Figure S8, Table S9). The likelihood was defined using a negative binomial distribution with mean defined by a cell type’s reference profile plus expected background and with a size parameter set at 10 to allow for extensive overdispersion. Reference profiles of immune and stromal cell types were acquired from previous work (41). New reference profiles were defined with the mean expression profiles of PanCK^+^ clusters in the UMAP projection. These new clusters were all PanCK^+^ and included five tumor-specific clusters and one cluster shared across all tumors. The tumor-specific clusters were labeled “tumor” and the shared cluster was labeled “epithelial”, and this interpretation was confirmed by a pathologist review.

The matrix of single-cell gene expression profiles was analyzed using UMAP, and each cell was assigned to a cell type as described above (Figure 3A). It should be emphasized that for classic single-cell studies, this UMAP-Cell-Type combination plot represents the essential information content of the experiment. For spatially resolved studies, however, this basic data type is the very beginning of a rich spatial analysis. In spatial profiling, each cell in this UMAP-Cell-ID representation has additional information of high-resolution X, Y, and Z spatial coordinates associated with it. For example, we observed that in these tumors, B cells typically gathered in dense clusters accompanied by T cells (Figure 3B). Plasmablasts also gathered densely, often proximal to smaller numbers of T cells. Macrophages both gathered in small clusters and trafficked diffusely throughout tumors. Neutrophils were usually found filling large vacancies within tumors, accompanied by very few other immune cells (Figure 3B).

**Figure 3.**
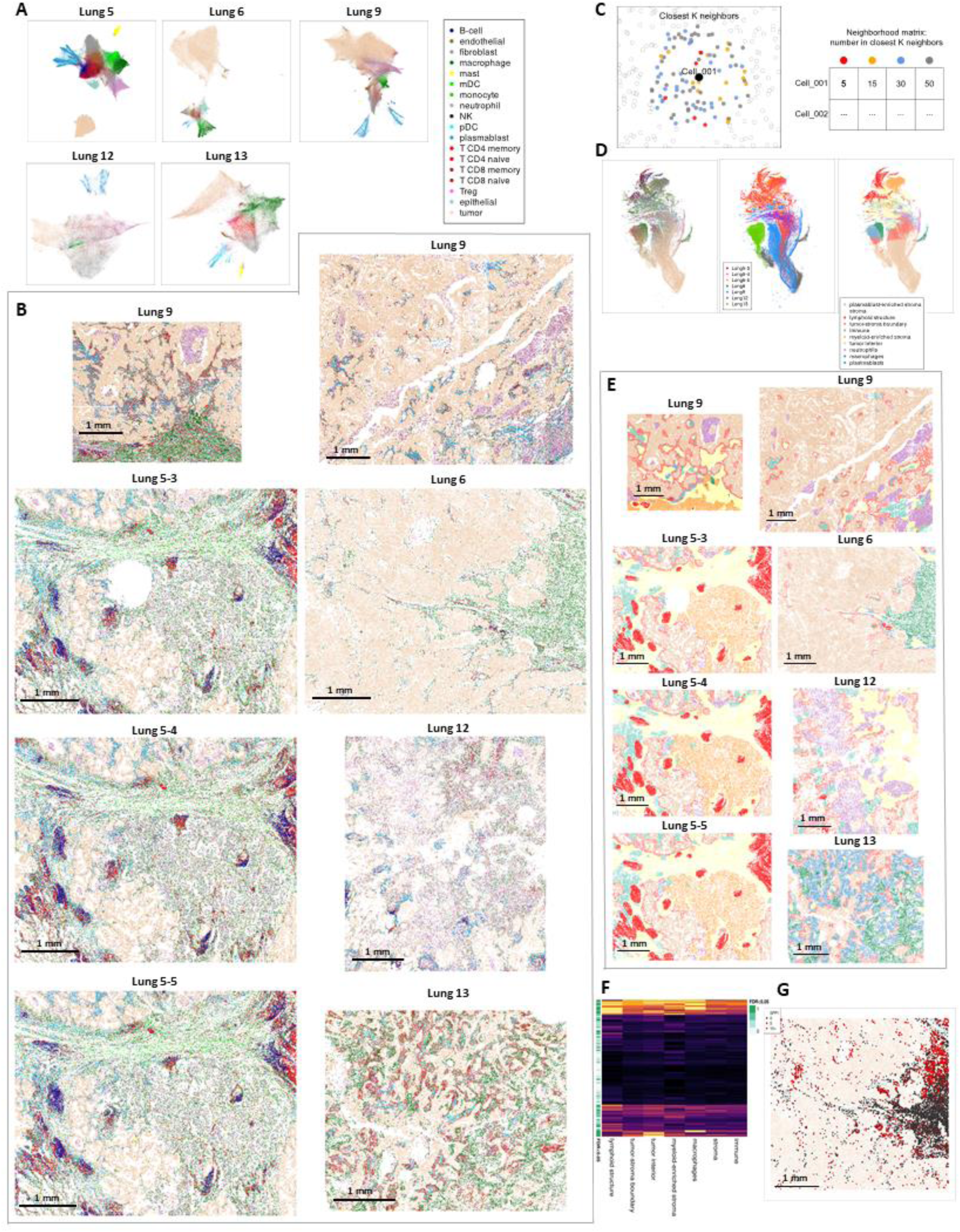
Spatial RNA Detection to Identify Cell Types and Cell-Cell Interactions in FFPE Human NSCLC Tissue. **A.** UMAP projections of each tissue based on the gene expression matrix. Color denotes cell type. **B.** Cells in physical locations (X, Y coordinates). Color denotes cell type. **C.** Definition of a neighborhood matrix. For each cell, the nearest neighbors are identified, and a summary of those neighbors is recorded. Here, the abundance of each cell type is used. This operation is performed for all cells, defining a matrix of cells and neighborhood characteristics. **D.** UMAP projection and clustering of cells based on the neighborhood matrix. Left: colored by cell type. Center: colored by tissue. Right: colored by clustering of neighborhood data or “niche”. **E.** Spatial arrangement of niches. For each cell, the frequency of each cell type among its 200 closest neighbors is recorded. Cells are then clustered into “niches” based on this data. Cells are shown in their physical locations and colored by their niche. **F.** Mean gene expression of macrophages in each niche in Lung 6. The green sidebar shows statistical significance from a global likelihood ratio test for uniform expression across all niches. **G.** Macrophage expression of SPP1 in Lung 6. In this tissue, macrophages located inside the tumor express SPP1, as shown in the upper half of the dense macrophage region. In contrast, the macrophages located along the vasculature and lymphoid cells in the tumor rarely express SPP1. Macrophages are shown in bold points, SPP1 expression in macrophages is shown on a color scale of black to red. Scale bar: 1 mm. These data can be interactively viewed at higher resolution in the CosMx Data Viewer (https://nanostring.com/products/cosmx-spatial-molecular-imager/ffpe-dataset/).

This spatial information also allows detailed analysis of cell “neighborhoods”. We defined a “neighborhood matrix” encoding the number of each cell type among each cell’s 200 closest neighbors (Figure 3C). Neighborhood matrices can be tailored to answer a broad range of different biological questions. For example, neighborhoods could be defined over smaller or larger distances, or a neighborhood matrix could encode average gene expression profiles or average gene expression profiles within specific cell populations.

Once a neighborhood matrix was defined, it was subjected to traditional single-cell analyses. A UMAP projection of our neighborhood matrix shows the diverse microenvironment states within these tumors (Figure 3D). For example, it shows neighborhoods of almost pure tumor cells with very low levels of macrophage and T-cell infiltration. Other neighborhoods were dominated by single cell types, including macrophages, neutrophils, plasmablasts, and myeloid dendritic cells (mDCs). Other neighborhoods held distinct mixtures of immune populations, such as B cells with T cells, macrophages with T cells and plasmacytoid dendritic cells (pDCs), and macrophages with neutrophils and infrequent lymphoid cells. Some of these neighborhoods were specific to single tumors, while others were shared across tumors. Finally, by clustering this neighborhood matrix, we partitioned the tumor microenvironment into distinct niches. Plotting niches in physical space clarified the spatial organization within and the contrasts between these tumors (Figure 3E).

Studies across larger numbers of samples require sample-level summary statistics. Using the cell types derived from the gene expression matrix, we found these tissues to differ in the relative abundances of the immune cell population within each tumor (Figure S9A). Using the niches derived from the neighborhood matrix, we identified additional differences in the multi-cellular niches comprising their microenvironments (Figure S9B). Niche abundances expand the information beyond what cell-type abundances alone can provide. For instance, samples Lung 5 and Lung 6 have similar macrophage abundances (7% and 8% of cells, respectively) while only Lung 6 contains the macrophage-dominated niche (9% of cells).

To define more nuanced sample-level summaries, we scored each cell for the number of tumor cells among its 100 closest neighbors, a metric of how much it has invaded into the tumor instead of remaining confined to the stroma. Contrasting this invasiveness score across cell types and across tumors revealed differences within and between tumors (Figure S10). For example, in Lung 6, macrophages were primarily surrounded by non-tumor cells, while neutrophils were more likely to be surrounded by tumor cells.

By contrasting the gene expression matrix and the neighborhood matrix, we examined further advanced questions for every gene, cell type, and neighborhood characteristic: “How does this cell type change expression in response to this neighborhood characteristic?” and “How does this dependency vary across tissues?” To answer these questions, we investigated the changes of gene expression in macrophages between niches in Lung 6 (Figure 3F). More than 43% of genes (415 of 960 genes) had expression changes between niches with false discovery rates below 0.05. The most statistically significant gene was SPP1 (*p*-value of 5 x 10^-61^). SPP1 has been shown to mediate macrophage polarization and up-regulate PD-L1 expression (21). Plotting macrophage SPP1 expression across the physical space of Lung 6 demonstrated two clear subpopulations of macrophages (Figure 3G). SPP1-high macrophages dominate the tumor interior and the upper half of the stroma, whereas SPP1-negative macrophages dominate the lower half of the stroma and the long thrust of diverse immune cells cutting deep into the tumor along the vasculature. Spatial expression analysis of SPP1 and HLA-DQA1 transcripts together (Figure S11) revealed these genes to be expressed in mostly mutually exclusive regions of the tumor, suggesting an antigen-presentation role for the SPP1-negative macrophages. Plotting the density of macrophage SPP1 expression across tumors identified changes between tumors and between niches (Figure S12). For every tumor, macrophages in the diverse “immune” niche had lower SPP1 expression than in almost any other niches.

With the spatial map of cell types in place, we turned to interrogate the interactions between tumor cells and T cells. First, we annotated 100 canonical ligand-receptor (LR) partners within the 980-plex panel (Figure 4A). Within our panel, many LR pairs relevant to the tumor immune interface can be found, including various immune checkpoints such as PD-L1/PD-1 and CTLA4/CD86. To understand how these interactions are changing across space and between samples, we devised a novel computational method to search for coordinated LR expression in neighboring cell types (Figure 4B). Using this method, we discovered 16 LR pairs that were enriched at the tumor-T cell interface in at least one of five lung tumors (Figure 4C). Many of these interactions were present in only a subset of the tumors. PD-L1/PD-1 (CD74/PDCD1) exhibited a higher interaction score across Lungs 5, 9, 12, and 13, but remained lower in Lung 6. Notably, HER2 (ERBB2) shows a similar profile across tumors. However, a member of the same receptor family, EGFR, maintained a higher interaction score in Lung 6 and decreased interaction in all the other lung tumors. This between-tumor variability in LR-signaling strength is consistent with the known variability in tumor response to immune checkpoint inhibitors and EGFR inhibitors.

**Figure 4.**
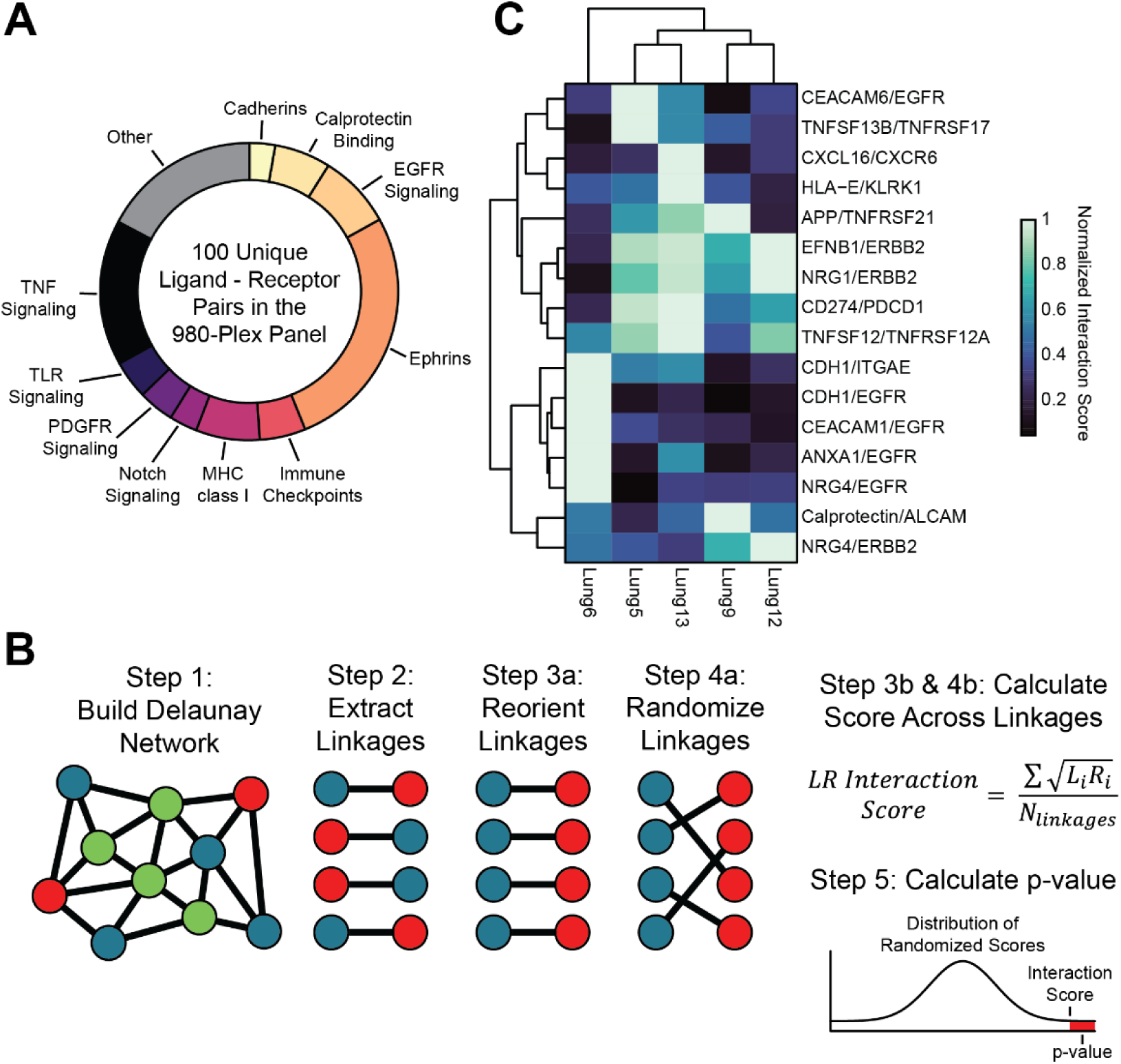
Paired Ligand-Receptor Expression Between Interacting Tumor and T Cells Varies Across Tumors. **A.** 100 unique ligand-receptor pairs are included in the SMI 980-plex panel. These interactions fall into functional categories in relative proportion. **B.** Ligand-receptor (LR) interactions are scored by first building a Delaunay network based on cellular spatial locations. The desired linkages are extracted from the network and used to calculate the LR interaction score, which measures pairwise LR expression between specified cell types. Linkages are reoriented to differentiate between ligand- and receptor-expressing cells. This score is compared to a distribution of scores produced using randomized linkages. Such a comparison can determine if the given configuration of interactions between specified cell types significantly enriches for pairwise LR expression. **C.** 16 ligand-receptor pairs exhibited spatial significance in at least one of five different lung cancer tumors. The interaction scores for these pairs are scaled, each score has a maximum of 1 across all tumors.

### 5. Reproducibility Across Serial Sections of FFPE NSCLC Tumor

To obtain a better understanding of the reproducibility of the SMI platform, three serial sections from an NSCLC tissue were profiled. Although these 3 serial sections would not contain the same individual cells, they had nearly identical tissue architecture (see Lung 5-3, Lung 5-4, Lung 5-5 in Figure 3), allowing comparisons of the same tissue region across the slides. This experiment offers an opportunity to observe technical variability that tests all aspects of the SMI platform: independent tissue preparation, cyclic chemistry, imaging, primary-secondary-tertiary data processing, and analysis in solid tissue with minimal biological variability.

As a first step, we examined the total-slide integrated RNA expression profile from each section by acquiring the total counts of each gene across all cells. These whole-slide integrated bulk profiles, each from 94,977 to 105,903 cells, were highly concordant. The lowest correlation between the log-scale expression profile of any pair of the three replicates was 0.989 (Figure S13).

To demonstrate reproducibility on a smaller spatial scale than a whole section, we partitioned two replicate (rep) sections into 74 grid squares each (Figure 5A, top). For this analysis, we used slides Lung 5 rep 3 and Lung 5 rep 5, which were the most spatially well-aligned pair. Grid squares contained between 600 and 2,000 cells; six grid squares intersecting a tissue-hole in rep 3 were discarded (Figure 5B). The total expression profile of each grid square was acquired to produce a gene expression matrix of 960 genes across 74 grid squares in each replicate. The correlation of 960-gene expression profiles between matching grid squares was high; the average square had a correlation of 0.96 between the log-transformed expression profiles of two replicates, and 95% of all squares had correlations between 0.87 and 0.94 (Figure 5B). The lowest correlation occurred in a square where the biological structure in serial sections differed; in Lung 5 rep 3, the square contained a small part of a tertiary lymphoid structure, while in Lung 5 rep 5 it did not.

**Figure 5.**
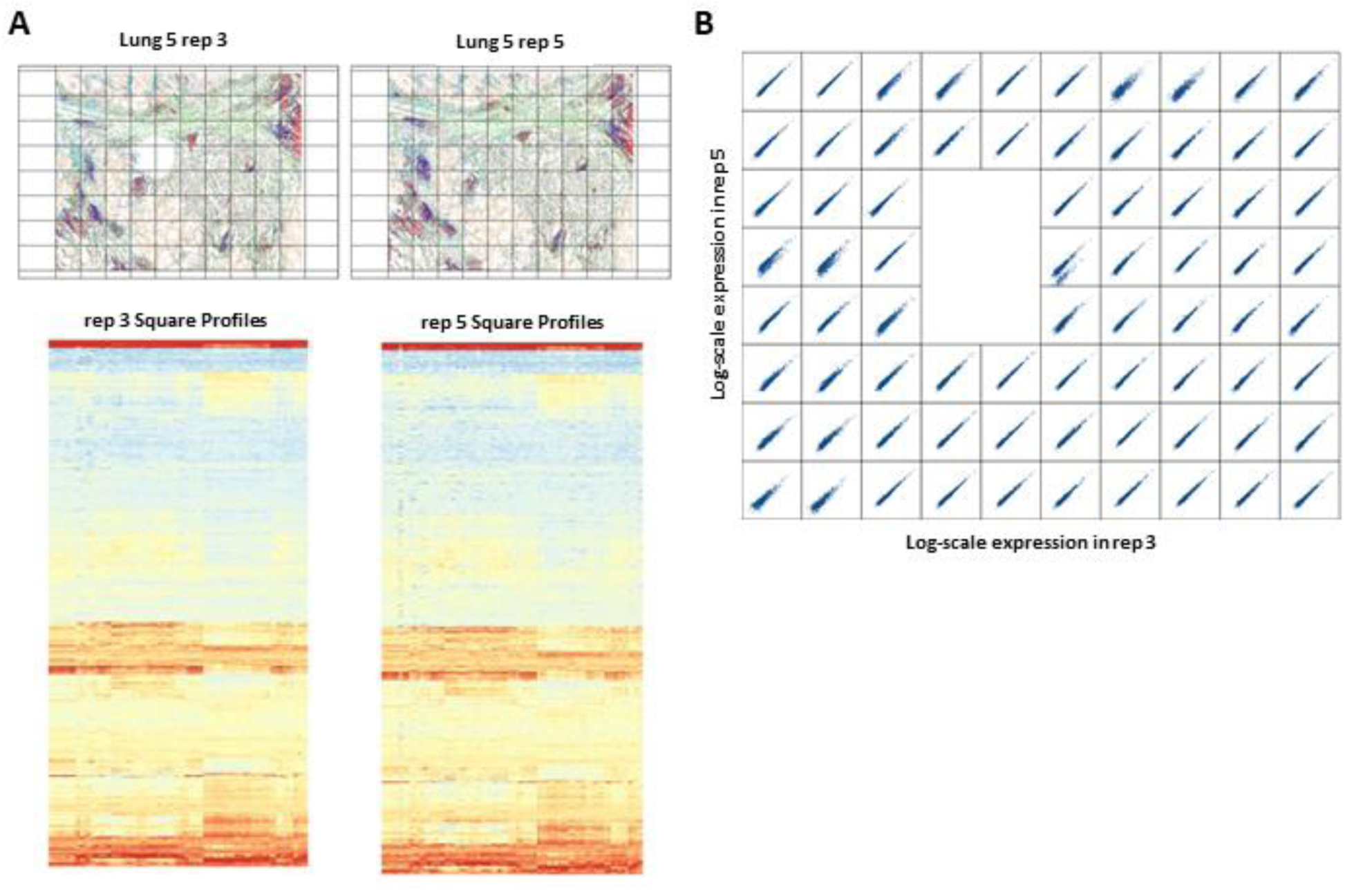
Concordance Between Serial FFPE Lung Sections Over a Spatial Grid. **A.** Each serial section of FFPE lung tissue (Lung 5 replicates 3 and 5) was partitioned into a grid. Squares held between 600 and 2,000 cells. **B.** Concordance between the 980-gene expression profiles of matching grid squares. Six squares overlapping a hole in rep 3 are excluded. To the right of the hole, a square with worse concordance is visible; this square contains part of a tertiary lymphoid structure in the rep 3 slide but not in the rep 5 slide.

### 6. High-Plex Protein Imaging using the SMI Chemistry

The SMI encoded detection chemistry utilized for RNA can be applied to protein detection by conjugating the oligonucleotide-readout sequences to antibodies (Figure S14). This simple expansion of the SMI chemistry for antibody-based detection is enabled by the extremely small size of the oligonucleotides required for SMI 64-bit encoding; only 60 to 80 nucleotides are required to encode all of the multiplexing.

For the 108-plex protein panel described here, a set of 129 barcodes with HD5 and HW4 were generated using the same initial set of 64-bit barcodes used for RNA. The barcodes were designed to maximize the bit distribution over 16 hybridization rounds and to minimize bit overlap across barcodes. A subset was selected to detect target proteins with the remaining barcodes as blank controls for misidentification quantification.

Using modified SMI oligonucleotide barcodes, we performed site-specific conjugation of 108 antibodies against immune cell activation states and drivers of cancer progression (Table S10). Among these, 104 antibodies carried a 64-bit encoded barcode to be read out over four detection events across 16 rounds of reporter hybridization. Four antibodies (PanCK, CD45, CD68, and membrane marker CD298) were conjugated to a distinct set of oligonucleotide landing sites for non-encoded morphological visualization during a single hybridization event with a distinct set of fluorescent reporter oligonucleotides. Each conjugated antibody had been previously reviewed by a pathologist and validated on GeoMx DSP (20) or via single-plex IHC.

FFPE breast cancer tissue of HER-2 positive invasive carcinoma was prepared under standard IHC conditions with an overnight antibody incubation. Following washes and fiducial application, the tissue sample in the flow cells was placed on the SMI instrument. Protein localization patterns (Figure 6A and 6B) and relative expression levels were read out on the SMI instrument using the same cyclic chemistry as the RNA readout assay. The primary data output is decoded OME-TIFF files, showing the localization pattern and relative intensity of each protein target in the assay (Figure 6).

**Figure 6.**
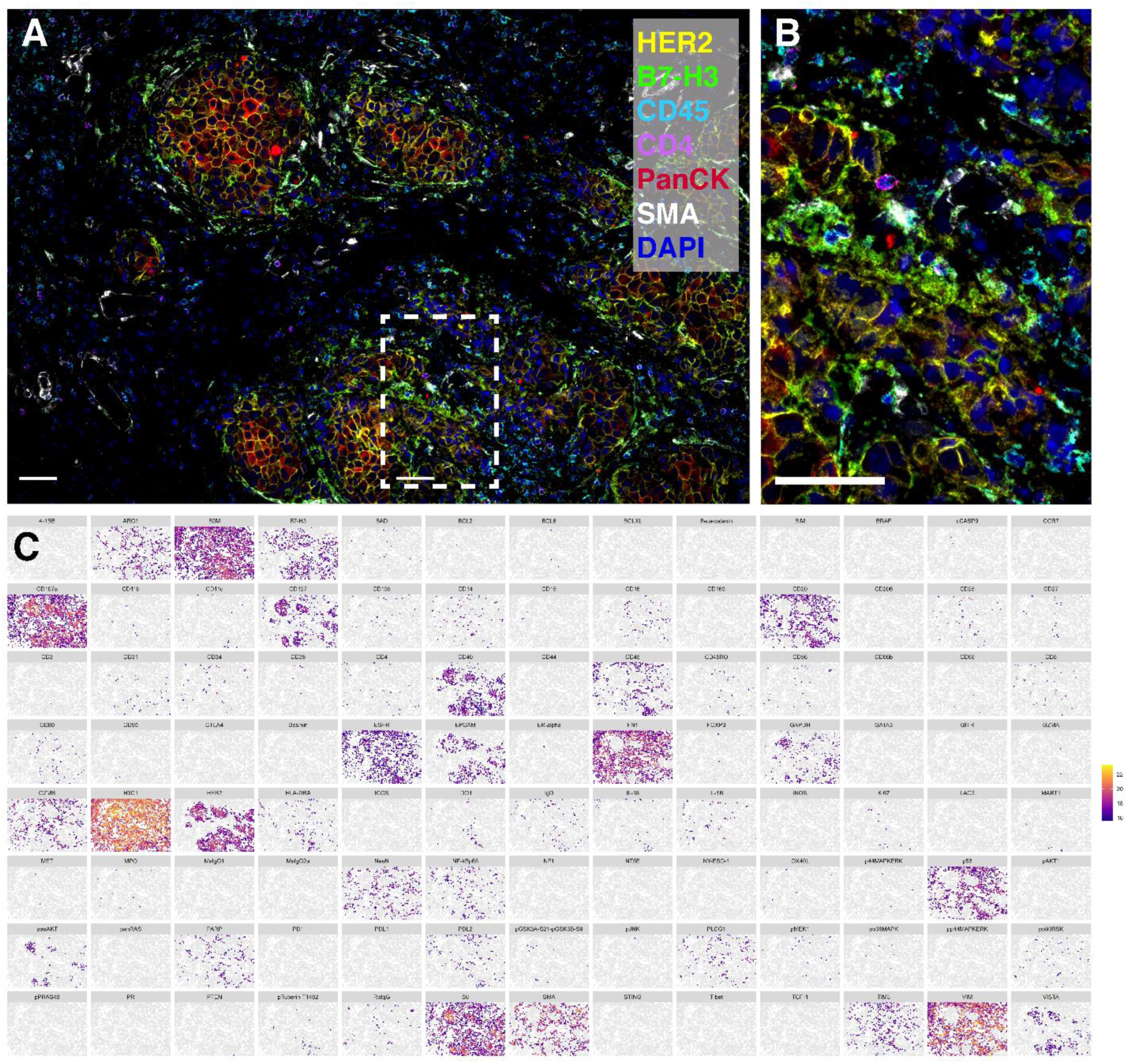
Spatial Subcellular Protein Analysis on SMI. **A.** Multi-channel overlay of six protein targets detected in a breast cancer biopsy (HER-2 positive invasive carcinoma) from a 108-plex assay (104 encoded targets and 4 morphological markers). **B.** Enlargement of boxed region in (A). Each decoded marker is visualized along with DAPI (blue) and the morphological markers HER2 (yellow), B7-H3 (light green), CD45 (cyan), CD4 (light purple), PanCK (red), and SMA (white). Scale bars: 50 µm. **C.** Single-cell expression profiles across 104-encoded protein targets in the sample shown in (A). Colored according to the log2 transformation of the sum of pixel intensities in each cell with a threshold for visualization the geometric mean of the negative isotype controls (mouse IgG1, mouse IgG2a, rabbit IgG) plus 1.5 standard deviations.

The OME-TIFF files were subset using cell segmentation masks to yield cell-by-cell expression patterns (Figure 6C, Figure S15). Protein localization patterns for a subset of targets, including CD45, Vimentin, CD3, and Histone H3, were cross-validated by comparing the IF signals from antibodies detected using traditional IHC or DAPI stain with a single target detected per channel to the computationally decoded protein maps (Figure S16). All other localization patterns were further validated by a pathologist’s review of decoded protein localization patterns using appropriate control tissues.

While this identical SMI chemistry has been successfully utilized to plex over 900 RNA-targets, the ability to multiplex proteins needed to be established. The plex-testing was accomplished by performing “nested-multiplexed” protein assays, across various plex levels on 35 FFPE cell lines. In a nested-multiplexed assay, we compare a 25-plex result with a 50-plex assay, where the 50-plex assay consists of the same targets as the 25-plex assay with 25 additional new targets. In this manner, the counts of the 25 targets measured in the 25-plex assay were graphed along the x-axis, and the abundance of the same 25-targets when measured as part of the 50-plex assay were graphed along the y-axis (Figure S17). The measured intensities should be independent of the plex of the final assay. The nested protein assays at 25-plex, 50-plex, and 79-plex yielded highly concordant results over all FFPE cell lines (Figure S17). These data suggest that protein detection on SMI can likely achieve well over 79-108 plex and may scale in a manner very similar to that of the RNA-based assay.

## Discussion

Real-world FFPE cancer samples are challenging to image at high-plex in a multiomic manner using conventional methods. As shown in Table S5, over 96% of the 800,327 total cells in the samples passed quality control and were analyzable at 980-plex RNA with subcellular spatial resolution. Especially noteworthy, this high efficiency of detection was independent of the overall RNA integrity of the samples. Samples Lung 5 and Lung 13 had QV200 values under 30% and are classified as “too degraded” to even attempt to perform bulk RNA-sequencing (42) while Lung 9 and Lung 12 would be graded as “medium” RNA quality. The fact that SMI had over 96% cell detection efficiency across this wide range of QV200 (and RIN scores) reflects the ability of the very small hybridization footprint of the SMI chemistry (35-50 nt hybridization zone) to quantitate very fragmented RNA patterns where the QV200 scores enter the “too degraded to sequence” classification.

The ultra-small readout zone also permits the very simple “migration” of the SMI chemistry from detecting *in-situ* RNAs to detecting oligo-labeled antibodies. Our 60-80 nt oligonucleotide-labeled antibodies are easily conjugated and purified and retain their original specificity and sensitivity. Over 80% of the antibodies in this SMI protein panel migrated directly from the extensively validated antibody panels developed for the GeoMx DSP (20). All of the DSP labeled antibodies successfully migrated to an SMI oligo-labeled antibody format. The remaining 20% were screened prior to and after conjugation in single-plex chromogenic assays and reviewed by a pathologist. To the best of our knowledge, this represents the highest plex protein imaging experiment carried out on FFPE samples.

Another unique feature of the SMI platform is the ability to work with FFPE samples and utilize standard, automatable FFPE sample preparation methods. SMI applies a sample preparation protocol nearly identical to standard low-plex *in-situ* hybridization, which does not require tissue clearing and/or expansion. Sample preparation time for an SMI measurement is a total of 1 day, with less than 60 min of hands-on time. All of these procedures can be automated on a standard tissue sample processor such as the Leica Bond system (43). The simplicity and automated features of the SMI sample preparation save time and labor and keep the tissue preparation highly consistent and reproducible (see Reproducibility Figure 5). The SMI chemistry uses a controlled number of 15-60 fluorescent dyes encoded into a single 64-bit imaging barcode, providing the sufficient fluorescence signal-to-noise ratio to quantitate a single *in-situ* molecule of RNA or protein even in the presence of severe autofluorescence of FFPE tissue samples.

SMI has demonstrated detection of up to 980-plex RNA and up to 108-plex protein, which is the highest plex of spatial RNA and protein profiling on FFPE tissue available to date with single-cell and subcellular resolution (Figures 1 and 6). Currently, RNA readout can be simultaneously performed with morphology marker antibodies, but not compatible with the 104-plex encoded protein panel yet. Increases in the plex of both RNA and protein assays are possible under this encoding scheme but have not yet been demonstrated. High accuracy is ensured by the error-robust color encoding scheme with a large Hamming distance (HD ≥ 4) between targets. In addition to the high sensitivity and specificity of RNA and protein detection, the very low cell drop-out rate (less than 8%, average of 4% in NSCLC samples) makes the data from almost every cell available for analysis, which has a distinct advantage over gene dropout in droplet-based single-cell sequencing methods (44).

SMI accomplishes RNA and protein multiplexing using a true 64-bit encoding method. This contrasts with techniques that are cycle non-encoded reagents for RNA and protein imaging. For example, cyclic IF methods (45) are limited to the number of fluorescent channels multiplied by the number of reagent imaging cycles, which increases linearly in the number of cycles of reagents (*e.g.*, 16 imaging rounds x 4 channel detection = 64-plex capability). SMI uses *encoded* barcoded antibodies and the same 16 imaging rounds with 4-channel detection, HD4 and HW4 yields a 1,210-plex capability. This class of encoded barcode technology provides the chemistry that can be essentially unlimited in plex with no fundamental change in instrumentation.

The multiomic capability of SMI is important for a number of key features. While high-plex RNA images supply incredible information content concerning the overall activity of individual cells, the images themselves do not reveal detailed tissue architecture as well as protein images; for instance, comparing a typical RNA tissue image (Figure 1C) with a typical protein tissue image (Figure 6A). The protein-based images clearly reveal overall tissue architecture in exquisite detail, whereas the RNA-image reveals “punctate-spots” distributed in a much more uniform pattern. SMI is capable to combine both the robust RNA information content with the tissue architecture in the image.

Simultaneous RNA and protein detection on a single slide is also required for accurate assignment of RNA to single cells, especially in real tissue materials, where cells are of very divergent sizes and densities. Cell segmentation methods relying on nuclear staining to predict cell boundaries are particularly challenged by low- and high-density tissue cell environments. SMI uses nuclear stain and antibody-based universal membrane stain cocktails, along with state-of-the-art cell segmentation algorithms, to accurately outline the cell boundary. This is essential for high-quality single-cell data which enables downstream analysis including cell typing, cell state, and cell-cell interactions (see Figures 3 and 4).

The spatial imaging data from five lung tumors provides more richness than can be fully explored in a single manuscript. For experts in immuno-oncology, the images in Figure 3 will suggest insights and questions far beyond those discussed here. We expect that similar datasets may often lead to multiple publications, as new analysts discover additional insights. We demonstrated a narrow application of spatial differential expression, and we looked for spatial dependency in a single cell type in a single sample. Using our data, similar analyses may still be performed for 959 other genes in any of 18 cell types and in any of the 5 tumors. These results could then be contrasted across samples. Gene expression could be contrasted with more nuanced spatial variables, such as the expression profile among cells’ neighbors or in the expression profile of only a cell’s neighbors of a given cell type. Experts in a given cell type or gene will find much to explore in these data.

With the advent of spatial molecular imaging using technologies such as the SMI platform, we can now directly test for the enrichment of pairwise ligand-receptor expression at the interface of interacting cells. Here, we find that out of unique ligand-receptor pairs, 16 were significantly enriched at the interface between tumor and T cells in at least one of five tumors. As expected, the interaction scores for each of these pairs varied across the tumors. This approach can be employed for any pair of cell types, and contrasts in ligand-receptor interactions can be drawn between different samples as well as between spatial contexts within a single sample.

We have not attempted a comprehensive analysis of all of the data in this first manuscript describing the SMI technology. For that reason, we are placing the raw and processed data described in this study into the public domain (http://nanostring.com/CosMx-dataset), where interested scientists from around the world can explore the data. Spatial imaging technologies have the potential to greatly enhance the field of Spatial Biology. When SMI is combined with high-plex profiling technologies (such as Digital Spatial Profiling), a truly comprehensive spatial investigation of tissues can be accomplished.

## Methods

### FFPE Cell Pellet Microarrays and Tumor Tissues

Custom FFPE cell pellet arrays of 16 cell lines for RNA assay and 35 cell lines for protein assay were made with A-FLX™ FFPE CELL PELLET by Acepix Bioscience, Inc. All cell lines were originally sourced from ATCC. The 16 cell lines for RNA assay include CCRF-CEM, COLO205, DU145, EKVX, HCT116, HL60, HOP92, HS578T, IGROV1, M14, MDA-MB-468, MOLT4, PC3, RPMI-8226, SKMEL2, SUDHL4. The 35 cell lines for protein assay include SHSY5Y CA, A431 CA, NCIH2228, DBTRG05MG, SW48, NB4, U251MG, U87MG, SUPB15, THP1, WSUNHL, SKMEL2, SKMEL5, SKBR3, RAMOS, RPMI8226, SUDHL6, RI1, RAJI, MDAMB468, HUT78, HUH7, HL60, HCT116, HCC78, H596, A431, HEK293 ICOS, HEK293 PD1-overexpressing, HEK293 CTLA4-overexpressing, HEK293 GITR-overexpressing, HEK293 LAG3-overexpressing, HEK293 PDL2-overexpressing, and HEK293 PDL1-overexpressing.

The breast cancer biopsy used for protein assay was CTRL301 tissue microarray supplied by US Biomax, Inc. The FFPE sample used was a malignant breast biopsy from a 61-year-old female with invasive carcinoma (Her-2 3+), T2N0M0 at IIA grade. Human tissues were collected under HIPPA approved protocols (US Biomax, Inc.).

Five FFPE human NSCLC tissues were acquired from ProteoGenex, Inc. All of these samples were collected under an ethics committee (European IRB analog) and with informed consent.

### SMI Instrument

SMI utilizes standard sample preparation methods typical for FISH or IHC on FFPE tissue sections, with the introduction of fluorescent bead-based fiducials that are fixed to the tissue to provide an optical reference for cyclic image acquisition and registration. For sample-processing details, see the “RNA assay FFPE tissue prep by manual method” section.

Following hybridization of ISH probes or antibody incubation, slides were washed and the coverslip with a defined height (50-100 μm) spacer was applied to the slide. The slide plus the coverslip constitutes the flow cell, which was placed within a fluidic manifold on the SMI instrument for analyte readout and morphological imaging.

The *in-situ* chemistry and imaging analyses were performed on a prototype instrument with a custom large FOV, high numerical aperture, and low aberration objective and associated optics that permit more than 6,000 cells to be imaged per FOV from a flow cell assembled onto a standard glass slide. The prototype system can run up to 4 flow cells simultaneously and can interleave fluidics and optical-imaging operations to maximize throughput.

The optical system has an epi-fluorescent configuration that is based on a custom water objective with a numerical aperture (NA) of 0.82 and a magnification of 13X. The FOV size was customized to 0.7 mm x 0.9 mm. Illumination is widefield with a mix of lasers and LEDs (130 W/cm^2^ at 488 nm, 69 W/cm^2^ at 530 nm, 26 W/cm^2^ at 590 nm, 81 W/cm^2^ at 647 nm) that allow imaging of Alexa Fluor-488, Atto 532, Dyomics Dy-605, and Alexa Fluor-647 as well as cleaving of photolabile-dye components. The camera used is FLIR BFS-U3_200S6M-C based on the IMX183 Sony industrial CMOS sensor, and the sampling at the image plane is 180 nm/pixel. An XY stage moves the flow cell above the objective lens, and a Z-axis motor moves the objective lens.

The fluidic system uses a custom interface to draw reagents through a flow cell using a syringe pump. Reagent selection is controlled by a shear valve (Idex Health & Science). A flow sensor between the flow cell and the syringe pump was used for flow rate feedback (Sensirion AG). The fluidic interface includes a flat aluminum plate that is in direct contact with the flow cell. The temperature of this metal plate was controlled to regulate the reporter hybridization temperature between 20°C to 35°C. The enclosure around the instrument was also maintained at a constant temperature using a separate thermoelectric cooler.

The CosMx SMI platform has been used for research use only to date.

### SMI *In-Situ* Hybridization (ISH) Probe Design

The ISH probes were designed to bind *in-situ* mRNA targets (Figure 1A). From 5′ to 3′, they each comprised a 35-50 nt target-complementary sequence followed by four consecutive 10-20 nt readout sequences corresponding to four “on” bits (HW4) assigned to each target. The target-binding sequences in ISH probes were developed by a probe design pipeline that optimizes sensitivity and specificity for mRNA transcripts. The process begins with an exhaustive evaluation of all possible contiguous 35-50 nt sequence windows for each mRNA target. This large pool of possible probe candidates was first filtered for ideal intrinsic characteristics including melting temperature (Tm), GC content, secondary structure, and runs of polynucleotides. Probes satisfying these parameters were further screened for homology to the full transcriptome of the parent organism utilizing the Basic Local Alignment Search Tool (BLAST) from the National Center for Biotechnology Information (NCBI). Preference was given to probes covering known protein-coding transcripts, lying within coding regions, and maximizing the coverage of the isoform repertoire. Final panel candidates were further screened for intermolecular interactions with other probes in the candidate pool including potential probe-probe hybridization as well as minimizing common sequences between probes. Five oligonucleotide RNA detection probes were designed per target mRNA. Negative control probes were modeled after synthetic sequences from the External RNA Controls Consortium (ERCC) set (34). These negative ISH probes were designed to contain the same intrinsic characteristics and subjected to the same inter-/intra-molecular interaction screens as the primary panel of probes.

The 10 to 20 nt readout sequences based on the 64-bit barcode design were filtered based on having a GC fraction greater than 35% to minimize the cross-hybridization between reporter probes and the junctions between readout sequences, as well as to maintain a Hamming distance of four between sequences. Also, readout sequences contained only bases A, C, and T to maximize binding kinetics between reporters and ISH probes.

### SMI Reporter Design and Assembly

The SMI reporter is a defined 15-to-60-dye DNA construct assembled from three oligonucleotide motifs: dye (RPD), sub-branch (RPU), and nanoBarCode (nBC) (Figure S18, Table S11). All oligos were made from DNA amidite (ChemGenes), while the RPU and nBC also contain PC-linker for photocleavage. The photocleavable linker moiety is introduced to the GeoMx probes *via* the conventional phosphoramidite method of DNA chain assembly using a custom 2-cyanoethyl phosphoramidite reagent derived from 5-methyl-2-nitrobenzoic acid starting material in a 10-step synthesis (46). Commercially available photocleavage phosphoramidites, such as PC-Spacer (P/N 10-4913) and PC-Linker (P/N 10-4920) from Glen Research would accomplish the same function as this custom phosphoramidite reagent. The RPD contains 15 nt with a 5’-amino modifier used to conjugate Alexa Fluor-488, ATTO 532, Dyomics-605, or Alexa Fluor-647 fluorophores. The RPU contains a repeated 10-20 nt nBC motif on the 5’ end, followed by a PC-linker and a repeated 10-20-nucleotide RPD motif toward the 3’ end. The nBC contains a 10-20 nt ISH-probe motif at the 3’ end, followed by a PC-linker and a repeated RPU motif at the 5’ end. Each RPD has one corresponding RPU and 16 nBCs for a total of 64 reporters.

Lyophilized RPD and RPUs were normalized to 1.5 mM and 0.5 mM in Tris EDTA (TE) pH 8, respectively. The nBCs were normalized to 50 μM in TE. All oligos were stored at 4°C prior to use. Reporter assembly occurs via a two-stage process involving RPD-RPU hybridization and nBC hybridization (Figure S18). The design of the “N” oligo regions (14-mers, Table S11) utilizes the observation noted by Zhang *et al.* (47) that sequences composed of 3 (of 4 possible) nucleotides hybridize more rapidly than those with 4 of 4. Thus, we constructed all combinations of 14-mer sequences composed of ATG nucleotides and filtered for single-base repeats (no single-base repeats of > 4 nt), particularly poly-G (no repeats of G > 3 nt), at least 5 G bases, Tm ≥ 40°C (calculated for 5 nM oligo concentration, 750 mM NaCl using the SantaLucia method), and dG > 0 at 5°C. Any passing sequences were screened for inter-sequence compatibility for both hamming distance (≥ 4) and cross-hybridization (no cross-hybridization with any other sequence with a Tm > 32°C (based on thermodynamics of cross-hybridization using DNA folding/ hybridization algorithms and calculated for 250 nM oligo concentration, 750 mM NaCl using the SantaLucia method). The 980-plex RNA and 108-plex protein readout requires 16 sets of pools. Each set contains four reporters with four different fluorophores. The pools were made by diluting the 1 μM stock to a 5 nM/probe solution in 8.75X SSPE, 0.5% Tween-20 (%w/v) and 0.1% ProClin^TM^ 950. Pool identity and cleavage were assessed using biotinylated targets on a streptavidin-covered slide (Schott). Pools were stored at 4°C until ready to use.

### Antibody Conjugation

All antibodies were sourced from vendors in a BSA- and glycerol-free format. Antibodies were quantified using UV spectrophotometry and quality-checked using gel electrophoresis. Antibody heavy chains were prepared for conjugation using SiteClick™ azide modification (Invitrogen) of carbohydrate domains. Amine-modified oligo tags were conjugated to antibodies using modifications of Site-Click chemistry (Invitrogen) and heterobifunctional DBCO linker (Click Chemistry Tools). The resulting conjugates were HPLC purified and normalized to 200 μg/mL as described (20).

For the protein detection assay, the oligonucleotides resembled those used for the SMI RNA detection in that there were landing sites for four of the 64 nano-barcode reporters, corresponding to the 64-bit barcode (Figure S14). For imaging of tissue morphology and cell boundaries, the antibodies were conjugated to an oligonucleotide possessing a single landing site for one of four morphology reporters, which have the same overall structure as the nano-barcode reporters but bind a unique set of four sequences that are orthogonal to the 64 landing sites used in the readout assay. This allows the signals from antibodies used for morphological visualization to be eliminated prior to the high-plex RNA and protein readouts.

### SMI 980-Plex RNA Panel Design

For comprehensive RNA profiling, we designed a 980-plex panel (Table S1 and Figure S5E) to investigate the biology of single cells across tumors and diverse organs. To select panel content, 749 genes were selected to capture critical cell states and cell-cell interactions. The remaining genes were selected to optimize the panel’s power to distinguish between different cell types.

While cell state is a broad term encompassing a wide variety of cellular phenotypes and processes, we focused our curation on core pathways or environmental factors that are important across broad areas of physiology and disease. These included immune cell states, basic cellular processes (*e.g.,* apoptosis, autophagy), cellular structures that integrate environmental cues (*e.g.*, cytoskeleton, extracellular matrix), and stress or damage responses (*e.g.*, hypoxia, wound healing, DNA damage response) (Figure S5). Genes for these cell states were primarily curated from the literature, using hallmark genes for each set. Genes for cellular signaling pathways, including ligands and receptors, were curated from the literature, HUGO gene families, and the KEGG BRITE Ontology. Finally, target genes for common cell signaling pathways were included when those transcriptional outputs were known with confidence. Consensus target genes were determined for the MAPK (48), NF-κB (48–51), Interferon (52), Wnt (53–60), and Hedgehog pathways (61).

Cell state and cell signaling categories were also subjected to a data-driven expression level assessment. We employed 76 individual Human Cell Landscape Datasets to score for expression level across cell types and tissues. We removed genes that were highly unlikely to be detected in any tissue types. However, we utilized subject matter expertise to selectively retain genes of high biological interest, for example, T-cell checkpoint genes. While these genes may be rarely detected in healthy tissue, they are crucial to understanding the immune system’s role in disease (Figure S5).

Once genes of biological interest were selected, the remainder of the panel was chosen to inform cell typing. First, literature-driven searches identified 56 highly informative immune cell markers (41, 62, 63) and 6 adipocyte markers (64–67). The remaining genes were then chosen in a data-driven manner to maximize the contrast between different cell type’s expression profiles.

Specifically, for a given pair of cell types, we scored each gene’s “pairwise distance” as (|x1 – x2| / sqrt(x1 + x2)) max(x1, x2), where x1 and x2 are the gene’s expression levels in the two cell types. The left side of this equation was motivated by the *t*-statistic and the mean-variance relationship of the Poisson distribution. The right side was motivated by the assumption that every transcript of a gene offers a fixed amount of evidence for one cell type versus another, and that the total evidence from a gene will therefore vary in proportion to its total transcripts. While the exact statistical power conferred by a gene depends on the cell typing method is used, the above statistic formalizes the intuition that useful cell-typing genes will have two properties: they will vary strongly across cell types, and they will have high expression levels in at least one cell type.

For every gene, the pairwise distance between all pairs of cell types within each dataset was calculated. Then, the total pairwise distance between all cell types within each dataset was calculated over the genes already chosen for the panel. Cell typing genes were then chosen in a greedy manner to improve the pairwise distances between the least distant cell types across the datasets. To initially choose genes that were informative between many pairs of cell types, the first genes were chosen to maximally increase the 40^th^ percentile of between-cell distances. Over 224 successive iterations, this percentile was dropped to 0.005 to focus on increasing the distances between the most similar cell types.

### RNA Assay FFPE Tissue Prep by Manual Method

Five-micron tissue sections were cut from FFPE tissue blocks using a microtome, placed in a heated water bath, and adhered to Leica Bond Plus Microscope slides (Leica Biosystems). Slides were then dried at 37°C overnight and stored at 4°C.

To perform *in-situ* hybridization on the tissue sections, the tissue slides were baked in 37°C oven overnight and then baked in 60°C oven for at least 2 h. The tissue sections were dewaxed in xylene (Millipore) for 5 min twice, ethanol (Pharmco) for 2 min twice, and then the slides were baked in 60°C oven for 5 min. The tissue sections were subjected to the target retrieval step using RNAscope^TM^ Target Retrieval kit (ACD Bio) and heated at 100°C in a pressure cooker for 15-30 min. After target retrieval, the tissue sections were rinsed with diethyl pyrocarbonate (DEPC)-treated water (DEPC H_2_O) (Thermofisher), washed in ethanol for 3 min, and dried at room temperature for 30 min. On the dried slide, a hydrophobic barrier line was drawn around the tissue section using ImmEdge^TM^ Pen (Vector).

The tissue was then digested with Protease Plus (ACD Bio) spiked with proteinase K (ThermoFisher), ranging from 1 μg/mL to 5 μg/mL depending on tissue type, at 40°C for 15-30 min. The tissue sections were rinsed with DEPC H_2_O twice, incubated in 1:400 diluted fiducials (Bangs Laboratories) in 2X saline sodium citrate and Tween (SSCT; 0.001% Tween-20, Teknova) for 5 min at room temperature, and washed with 1X phosphate-buffered saline (PBS; ThermoFisher) for 5 min. After digestion and fiducial placement, the tissue was fixed with 10% neutral-buffered formalin (NBF) for 1 min to maintain soft tissue morphology, then washed twice with Tris-glycine buffer (0.1M glycine [Sigma], 0.1M Tris-base [FisherScientific] in DEPC H_2_O) for 5 min, and washed with 1X PBS for 5 min. The fixed tissue was blocked using 100 mM *N*-succinimidyl (acetylthio) acetate (NHS-acetate; ThermoFisher) diluted in NHS-acetate buffer (0.1M NaP+0.1% Tween PH8 in DEPC H2O) for 15 min at room temperature and washed in 2X saline sodium citrate (SSC) for 5 min. Adhesive SecureSeal^TM^ Hybridization Chamber (Grace Bio-Labs) was placed to cover the tissue.

NanoString ISH probes were prepared by incubation at 95°C for 2 min and immediately transferred to ice. The ISH probe mix (1 nM ISH probe, 40% formamide, 2.5% dextran sulfate, 0.2% BSA, 100 μg/mL salmon sperm DNA, 2X SSC, 0.1 U/μL SUPERase•In™ [Thermofisher] in DEPC H_2_O) was then pipetted into the chamber, and adhesives supplied with the chambers were applied to the ports of the chambers. Hybridization occurs at 37°C overnight after sealing the chamber to prevent evaporation. After ISH probe hybridization, the tissue sections were washed twice in a buffer composed of 50% formamide (VWR) in 2X SSC at 37°C for 25 min, washed twice with 2X SSC for 2 min each at room temperature, and then blocked with 100 mM NHS-acetate for 15 min. After blocking, the hydrophobic barrier was removed with a blade, and a custom-made flow cell was attached to the slide.

### Automated RNA Assay FFPE Tissue Prep on Leica Bond RX

As in the manual FFPE tissue prep method, tissue slides were baked overnight at 60°C to ensure tissue adherence to the positively charged glass slides (Leica Bond Plus Microscope slides, Leica Biosystems). Then the tissue was deparaffinized and digested with Protease Plus (ACD Bio) spiked with proteinase K (ThermoFisher) ranging from 1 μg/mL to 5 μg/mL depending on tissue type and prepared for heat-induced epitope retrieval (HIER) on a Leica Biosystems automated tissue handler (Bond RX, Leica Biosystems). For cell pellet array (CPA) or tissue microarray (TMA) samples, HIER requires the treatment in Leica buffer ER1 at 100°C for 8 min, while 30min treatment for tissue samples. After Leica handling, the remaining process is the same as RNA assay FFPE tissue prep by the manual method described above.

### RNA Isolation

Total lung RNA was isolated from single or double 20 μm FFPE curls using RNeasy FFPE Kit (Qiagen) and digested with proteinase K for 30 min. Lung RNA was quantified using an Agilent RNA 6000 Nano Kit (Agilent) according to the manufacturer’s instructions. DV200 and RIN scores were determined using Agilent BioAnalyzer 2100 Expert Software B.02.09. For DV200, Region 1 selection was from 200 nt to about 8,000 nt using Smear Analysis.

### Automated Protein Assay Sample Preparation on Leica Bond RX

Deparaffinization and antigen retrieval were performed automatically on a BOND RX machine (Leica Biosystems) using Bond Dewax Solution (30 seconds at 72°C) and Bond ER Solution 1 (20 minutes at 100°C). Samples were blocked for 1 hour with blocking buffer W (NanoString). Oligonucleotide-conjugated primary antibodies were pooled and diluted uniformly to 100 ng/ml in blocking buffer. Samples were incubated with primary antibodies overnight at 4°C. Then, samples were incubated with fiducials (0.00025%, Bangs Laboratories) for 5 minutes. To secure antibodies and fiducials, the samples were fixed with 4% paraformaldehyde for 10 minutes at room temperature and then washed in 1 x PBS. Prior to cyclic encoded protein detection, slides were incubated with *N*-succinimidyl (acetylthio)acetate (NHS-acetate, Thermo Scientific, diluted in 0.0932 M Na2HPO4 + 6.8 mM NaH_2_PO_4_ + 0.1% Tween pH 8) for 15 minutes on the SMI instrument.

### Cyclic RNA Readout on the SMI Instrument

Processed tissue was assembled into the flow cell and loaded onto the SMI instrument. The tissue was washed with 1.5 mL of reporter wash buffer to remove air bubbles in the flow cell. The Reporter Wash Buffer consisted of 1X SSPE, 0.5% Tween 20, 0.1 U/μL SUPERase•In RNase Inhibitor (20 U/μL), 1:1000 Proclin 950 and DEPC-treated water. A low-resolution image of the whole slide was acquired to allow the user to access the tissue, and then FOVs were placed at the areas of interest.

To start the cyclic RNA readout, 100 μL Reporter 1 was flowed in at 200 μL/min and incubated for 15 minutes. After incubation, 1 mL of Reporter Wash Buffer was flowed into the flow cell at 750 μL/min to wash out the reporter probes that did not hybridize. Following reporter wash, 100 μL imaging buffer was flowed into the flow cell prior to imaging. Imaging buffer consisted of 80 mM glucose, 0.6 U/mL of pyranose oxidase from Coriolus sp., 18 U/mL of catalase from bovine liver, 1:1000 Proclin 950, 500 mM Tris-HCI Buffer pH 7.5, 150 mM sodium chloride, and 0.1% Tween 20 in DEPC-treated water. After eight Z-stack images (800 nm step size) of each FOV were acquired, fluorophores on the reporter were UV cleaved (385 nm, 116 J/cm^2^, 500 ms) and washed off with 200 μl strip wash buffer. The strip wash buffer consisted of 0.0033X SSPE, 0.5% Tween 20, and 1:1000 Proclin 950. This fluidic and imaging procedure was repeated for the 16 reporter pools, and the 16-round of reporter hybridization can be repeated multiple times for increased sensitivity.

After all cycles were completed, the tissue was subjected to morphology stain workflow on the same instrument. The blocking buffer (Buffer W, NanoString) was incubated in the flow cell for 30 min, then the tissue was incubated with a 4-antibody cocktail (CD298, CD45, CD3, PanCK) diluted in Blocking Buffer for 1 h. After the incubation, the tissue was washed with 8 mL of Reporter wash and then 100 μL of imaging buffer was flowed into the flow cell, prior to collecting the antibody-labeled images of the tissue.

### Cyclic Protein Readout on the SMI Instrument

The on-instrument SMI protein assay readout was performed as described for RNA, with three exceptions. The readout was only 16 rounds of hybridization with no repeated cycling. Morphology visualization was performed using oligonucleotide-conjugated antibodies (Figure S14), as described in the Antibody Conjugation section. Tissue morphology and cell membrane morphology were visualized following hybridization of a specific pool of nano-barcode reporters to the oligonucleotide-conjugated antibodies.

### Primary Data Processing

Primary data processing is a standard image analysis of a three-dimensional multi-channel image stack obtained at each FOV location. The objective of this analysis is to reduce the multi-dimensional image stacks to a single list of individual reporters seen in the specific binding event. This process was performed in parallel across all FOVs and occurs in line with data acquisition. The image processing comprises three main steps – registration, features detection, and localization.

3D rigid image registration was performed with the use of fiducial markers embedded within the tissue sample. The fixed image reference was established at the start of the experiment prior to reporter hybridization. Subsequent image stacks, shifted by stage motion, were matched to this reference using phase correlation. Individual channels within the image were aligned to each other through the application of a pre-calibrated affine transformation.

The RNA image analysis pipeline focuses on identifying diffraction-limited features that represent the fluorescence response from a single molecule. Once the image stacks were registered, a 2D Laplacian of Gaussian (LoG) filter was applied to each channel to remove background and enhance the encoded reporter signatures. The kernel size and standard deviation of the filter were matched to the expected reporter point spread function. Post filtering, potential reporter locations were identified as local maxima using a 3D nearest neighbor search. Only local maxima greater than a channel-specific threshold were retained for further localization. Thresholds for each channel were predetermined empirically based on the signal-to-noise ratio (SNR) of the fluorescent channel. The retained maxima were assigned a confidence value determined by the intensity of the reporter signal.

Sub-pixel localization of each feature was obtained by fitting a 2D polynomial to the maxima and analytically solved for the sub-pixel maxima locations in X and Y. The final reporter signature locations in X, Y, and Z axes along with the assigned confidence were recorded in a list assigned to the specific reporter-binding event. After the acquisition was completed, all features pertaining to a given FOV were collated into a single list that forms the basis of the secondary analysis.

Protein image analysis pipeline requires a different analysis approach because, in general, multiple proteins have signals that overlap on a single pixel. The protein image analysis pipeline uses locally adaptive thresholds (68) to segment regions that range from single diffraction-limited features to large contiguous clusters within the FOV. Instead of quantification of proteins as individual diffraction-limited spots, we analyze regions of expression by linear unmixing of overlapping expression patterns combined with hard-decoding of thresholded encoding images. The protein is deemed potentially present if a pixel matches the expected encoding “on bits” within the 64-bit encoding scheme. From there, the protein intensity is computed by linear unmixing using pseudo inverse of the protein signature matrix. The output of the primary data analysis is a series of 2D binary maps that represent encoded reporter signals in each hybridization round. The binary segments are binned together based on encoded bits. The common regions between the binned masks generate protein localization masks. The localization masks for each protein are stored followed by linear unmixing using pseudo inverse of the protein signature matrix (PSM). The method used for pseudo inverse is Cholesky decomposition (69). The pseudo inverse of PSM is multiplied by the measured images, which provides protein expression and localization patterns. A more detailed description of the protein analysis pipeline will be the subject of a future manuscript.

### Secondary Analysis and Decoding

The imaging data converted to a table of XYZ locations of all individual reporter-binding events was used to determine the presence of individual transcripts. To start this process, each unique location with at least one reporter-binding event is considered a ‘seed’, and all neighboring locations to each seed with at least one reporter-binding event were determined. The neighbor search is limited to a radius of 0.5 pixels (90 nm) in a first pass through the data. Any seeds with fewer than 4 unique reporter probe-binding events in the neighborhood were removed from being considered for transcript decoding due to their inability to form a complete gene-specific barcode. All possible four reporter combinations of unique reporter probes in a seed’s neighborhood are then matched against a table of all potential gene-specific barcodes to detect the presence of a gene in a seed’s neighborhood. Only those four reporter combinations for which at least one of the reporter probes is detected at the seed location (as against the neighborhood around the seed) are considered valid for the target matching process. If more than one gene was detected in a seed’s neighborhood, the seed and all the transcripts detected in its neighborhood were dropped from further analysis.

All the seeds (or transcripts) retained after this step went through another filtering step. In this step, any seeds with a high probability of making a transcript call by random chance (and hence the transcripts detected in their neighborhoods) were then dropped from further analysis in the first pass through the data. Given that a set number of barcodes (*e.g.,* 980) out a pool of all possible four reporter barcodes that can be generated using 64 unique reporters (465, 920) were used to denote gene types, there is a non-zero probability that any random combination of four unique reporters can match a gene-specific barcode. This probability is further increased when the unique reporter binding events in a seed’s neighborhood can be used to generate many potential four reporter barcodes. To ensure high confidence in the presence of a true transcript for every target call made, any seeds (and associated transcript calls) with a random transcript call probability of more than 2% were dropped from further analysis in the first pass through the data. The reporter binding event locations that contributed to making transcript calls were used to estimate a centroid location for all transcripts retained after this filtering step. All reporter binding events that contributed to making retained transcript calls were then removed from the original imaging dataset and this modified dataset was used as a starting point for a second pass through the data.

The second pass through the data repeated all the steps above albeit with an increased target search radius of 0.75 pixels (135 nm). The rationale behind this increase in radius was to try and recover any potential transcript calls that may have been lost due to local tissue motion during reagent cycling. With the increased radius of 0.75 pixels, the transcripts added to the final list of transcript calls after adding a second pass through the data can vary from sample to sample and were found to range from approximately 20-40% of the total transcript calls in sample FOVs with single and dual radius analysis. After the conclusion of two passes through the data, we have a list of potential transcript locations in each FOV. However, as we consider each unique reporter binding event location as a ‘seed’ for transcript calling, there is a potential for duplicate calling of each individual transcript, artificially inflating the total count of transcripts detected in a FOV. For instance, if all four reporter binding events contributing to a transcript call occur at slightly different locations, the same transcript could be counted four times. To prevent this from happening, for each gene, all transcript calls made are filtered to ensure that there is no other transcript call present within a radius of 0.75 pixels from each transcript’s estimated location. Whenever multiple transcript calls are found within this search radius, the transcript call with the highest number of reporter binding events contributing to it is retained and others discarded.

### Cell Segmentation Algorithm

We established a cell segmentation pipeline combining image preprocessing and machine-learning techniques (Figure S3). Briefly, our pipeline takes tissue images stained with both nuclear and membrane markers (DAPI, CD298/PanCK/CD3) to perform rescaling, normalization, image deconvolution, and boundary enhancement. Image subtraction was performed between the nuclear and the membrane channels to enhance the contrast between adjacent cells while reducing the auto-fluorescence signal. The preprocessed images were fed into pre-trained Cellpose neural network models (70) for both nuclear and cytoplasm modes of segmentation. Results from two segmentation tasks were combined to select the best results from each mode by analyzing intersection and union between all segmented cells. The combination of the preprocessing steps, the dual-mode segmentation, and using the publicly available pre-trained models have greatly improved the robustness of cell segmentation outcomes, making retraining the models unnecessary for most tissue images.

The analysis pipeline has run with two different modes of hardware processors: Cuda GPU (Ge Force RTX 2070, Quadro P4000) and parallel CPU threads, with average segmentation runtime of around 94 secs per FOV with 5 channels, 6 to 8 Z-slices, and image size of 5,472 x 3,678 pixels. The final segmentation step was to map each transcript location in the registered image to the corresponding cell, as well as the cell compartment (nuclei, cytoplasm, membrane), where the transcript is located. Other features/properties generated include shape (area, aspect ratio) and intensity statistics (min, max, average) per cell.

### Segmented Regression to Estimate the Relationship Between SMI and RNA-seq

Segmented regression was performed using the R package “segmented”. Log-transformed SMI counts were modeled as a function of log-transformed RNA-seq counts. To avoid negative-infinity values after log-transformation, RNA-seq counts were thresholded below their lowest non-zero value. Estimated breakpoints and linear trends were extracted from the segmented regression models.

### Analysis of NSCLC Samples

Single-cell expression profiles were derived by counting the transcripts of each gene that fell within the area assigned to a cell by the segmentation algorithm. Cells with fewer than 20 total transcripts were omitted from the analysis. A normalized expression profile was defined for each cell by dividing its raw counts vector by its total counts. A separate UMAP projection was computed for each tissue.

After analyzing the single-cell expression data, we created a single-cell “neighborhood” matrix. This matrix specifies the number of each cell type among each cell’s 200 closest neighbors in 2D physical space. This matrix was then input into the UMAP algorithm. To define “niches”, this matrix was clustered using the Mclust algorithm.

To test whether each gene was differentially expressed between macrophages in different niches, a linear model was run predicting raw counts from the niche. Only the 6,884 macrophages found in Lung 6 were used in this analysis. A global *p*-value for each gene was taken using a likelihood ratio test comparing this linear model to the null model, using the R package lmtest. False discovery rates were calculated using the Benjamini Hochberg procedure via the R function p.adjust.

Following cell typing, we interrogated ligand and receptor signaling between adjacent cells. Delaunay triangulation was used to build a spatial adjacency network, given the spatial locations of each cell. For each pair of adjacent tumor cells and T-cells, we calculated an interaction score using the geometric mean of their ligand and receptor expression, respectively. This ligand-receptor interaction score was recalculated across all adjacent cell pairs for 100 distinct ligand-receptor interacting partners. Next, an average score was calculated for each ligand receptor pair. Finally, each average score was tested to determine if it was enriched by the spatial arrangement of cells within the adjacency matrix. This was performed by producing a null distribution of simulated average scores calculated using randomized adjacency networks.

### Protein Expression Visualization

The primary outputs of protein data processing are protein localization maps for each antibody in the assay, as well as images of the three antibodies used for tissue morphology and cell membrane visualization. Protein expression per cell was reported as a sum of the intensity values of pixels present in both the cell label and protein localization mask. A limit of detection for visualization was set by filtering to only cells with a detectable signal from at least one of the three isotype controls (19% of cells for Figure 6), using the geometric mean of the isotype control signals and adding 1.5 standard deviations. Isotype controls are non-specific negative control antibodies against rabbit IgG1, mouse IgG2a and mouse IgG1 (Figure S15). Visualization of protein localization patterns was performed in ImageJ. Single-cell expression visualization was performed in R.

## Data Availability

The full RNA NSCLC dataset used in this study is available at http://nanostring.com/CosMx-dataset.

## Code Availability

Data from this publication (the full RNA NSCLC dataset) has been put into the public domain in a format that can be analyzed and visualized using a variety of open-source packages, such as Seurat (https://github.com/satijalab/seurat) and Giotto (https://github.com/RubD/Giotto). Nearly all of the analyses performed in this paper can be accomplished using these open-source packages. For any of the specialized code/analyses performed in this manuscript that are not available through open-source packages, requests can be made through email to the corresponding author.

## Acknowledgment

Research and development reported in this publication was supported in part through a strategic development collaboration between NanoString Technologies and Lam Research (Fremont, CA). The authors thank Brian Birditt, Brian Filanoski, Justin Jenkins, and Edward Zhao from NanoString Technologies for providing technical support.

## Author Contributions

Shanshan He – Conception and design of the work, supervised data collection, analysis and Interpretation, and drafted the manuscript.

Ruchir Bhatt – Developed protein data processing algorithms.

Carl Brown – Instrument control software, SMI automation, integration of all instrument subsystems software.

Emily A. Brown – Reviewed and validated protein data (Figure 6), performed the nested protein multiplex validation experiment (Figure S17).

Derek L. Buhr – Development of manual and automated processes for inventorying, quantitation, normalization, pooling, purification, and quality control of oligonucleotides in 980-plex RNA panel and SMI reporters.

Kan Chantranuvatana – Performed oligo conjugations of antibodies used in SMI protein assays. Patrick Danaher – Designed the 980-plex RNA panel with EP, performed the comparison to RNA-seq in Figure 2, performed the NSCLC analyses of Figure 3, performed the reproducibility analysis of Figure 5, and manuscript writing/editing.

Dwayne Dunaway – Led team that developed the system, instrument, and software.

Ryan G. Garrison – Development of reporter chemistry, assembly, quality control and manufacturing. Contributed to the section “SMI Reporter Design and Assembly” and Figure S18.

Gary Geiss – Lead development of protein-based SMI assays, design and interpretation of protein experiments.

Mark T. Gregory – Analyzed the profiling data of human lung cells.

Margaret L. Hoang – Contributed to the development of reporters and analysis of FFPE RNA quality, created Figure S6.

Rustem Khafizov – Designed and developed reporter structure. Emily E. Killingbeck – Contributed to the SMI methods.

Dae Kim – Developed overall concept, designed and guided experiments, analyzed data. Tae Kyung Kim – Supervised data collection and interpretation of human lung samples.

Youngmi Kim – Supervised data collection, analysis, and interpretation of human lung samples.

Andrew Klock – Developed the SMI ISH probe design pipeline; designed SMI ISH probes used in 980-plex RNA panel.

Mithra Korukonda – Developed the primary data analysis pipeline for RNA and protein targets. Alecksandr Kutchma – Designed SMI ISH probes and wrote the section on probe design.

Zachary R. Lewis – Designed and developed the protein assay, executed protein analyses, manuscript writing and editing.

Yan Liang – Pathological review to identify the correct staining pattern of all antibodies used.

Jeffrey S. Nelson – My team has been responsible for all the chemistry process development efforts and supply-chain management pertaining to the outsourcing of oligonucleotide synthesis, process improvements, price negotiation and supply-agreement, key component scale-up and validation of all the custom reagents and DNA components in R&D that have been required for SMI – including CMO and validation of PC-spacer, synthesis and QC of the three large-scale component sets needed for SMI Reporter manufacturing plus high-throughput synthesis, outsourcing, QC and processing of the thousands of the required ISH probes.

Giang Ong – Developed morphology and segmentation markers.

Evan P. Perillo – Developed the SMI instrument optical subsystem, instrument validation and support.

Joseph Phan – Developed encoding scheme, screened reporter sequences, developed readout Sequences.

Tien Phan-Everson – Optimized protein assay, data collection, executed protein analyses, manuscript writing/editing.

Erin Piazza – Designed the 980-plex RNA panel with PD, manuscript writing/editing. Tushar Rane – Developed secondary analysis and target decoding pipeline for RNA targets, developed the SMI instrument fluidic subsystem and workflow software.

Zachary Reitz – Developed the ligand-receptor interaction analysis method, analyzed ligand-receptor interactions across all tumors, manuscript writing/editing.

Michael Rhodes – Initial chemistry development, supervised data releases, and manuscript writing.

Alyssa Rosenbloom – SMI Protein Assay content design and reagent validation.

David Ross – Created plots of transcript positions overlaid on morphology and segmentation images for Figures.

Hiromi Sato – Manuscript development, writing, and editing. Created figures and tables.

Aster W. Wardhani – Designed and developed the cell segmentation pipeline, manuscript writing/editing.

Corey A. Williams-Wietzikoski – Lung RNA isolation, processing, and quality measurement, NGS library preparation and sequencing.

Lidan Wu – Established cell segmentation pipeline, optimized the on-instrument SMI readout workflow, manuscript writing/editing.

Joseph M. Beechem – Conceived the project, helped in experimental design and analysis, manuscript writing/editing.

## Supplemental Materials

## Supplemental Tables

**Table S1. Genes for 980-plex RNA panel.**

**Table S2. Probe sequences and barcodes.**

**Table S3. 1,210 encoding barcodes for 980-plex RNA panel.**

**Table S4. CPA summary statistics.**

**Table S5. SMI results summary of lung tissue samples.**

**Table S6. Lung tissue sample information.**

**Table S7. RNA quality of lung samples.** Evaluation of RNA quality from FFPE samples (42).

**Table S8. Lung summary statistics.**

**Table S9. Expected expression profiles used in cell typing.** Each gene is scaled to have a maximum of 1.

**Table S10. Contents of 108-plex protein panel.**

**Table S11. Reporter readout probe sequences.**

---

**Figure S1.**
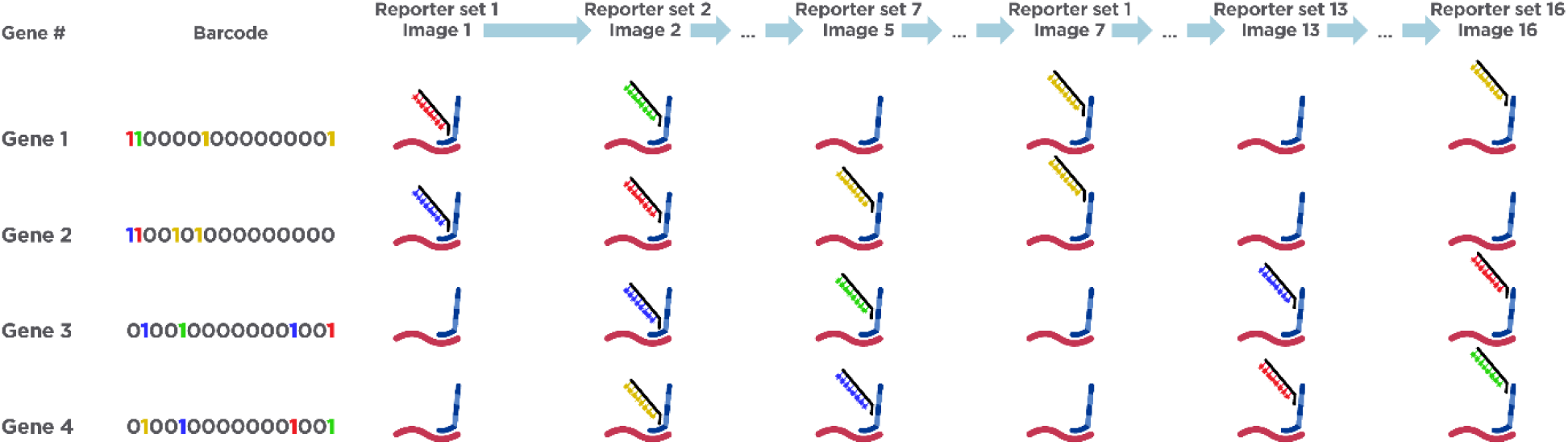
Schematic Depiction of SMI Encoding Design and Readout Process. Each gene is assigned with a unique 16-digit barcode with 4 “on” spots (labeled as “1”) and 12 “off” spots (labeled as “0”). Each digit of the barcode indicates the presence of the reporter that is associated with the target in the specific reporter hybridization round. “1” means that there is a reporter hybridizing to the ISH probe of the target in that hybridization round, and its color indicates the fluorophore of the hybridized reporter. “0” means that there is no reporter that binds to the target ISH probe in that hybridization round, and the target should be silenced or blank in that round of imaging. For each gene, 4 reporters will sequentially bind to the 4 designated reporter landing domains of the ISH probe throughout the 16 rounds of cyclic reporter readout.

**Figure S2.**
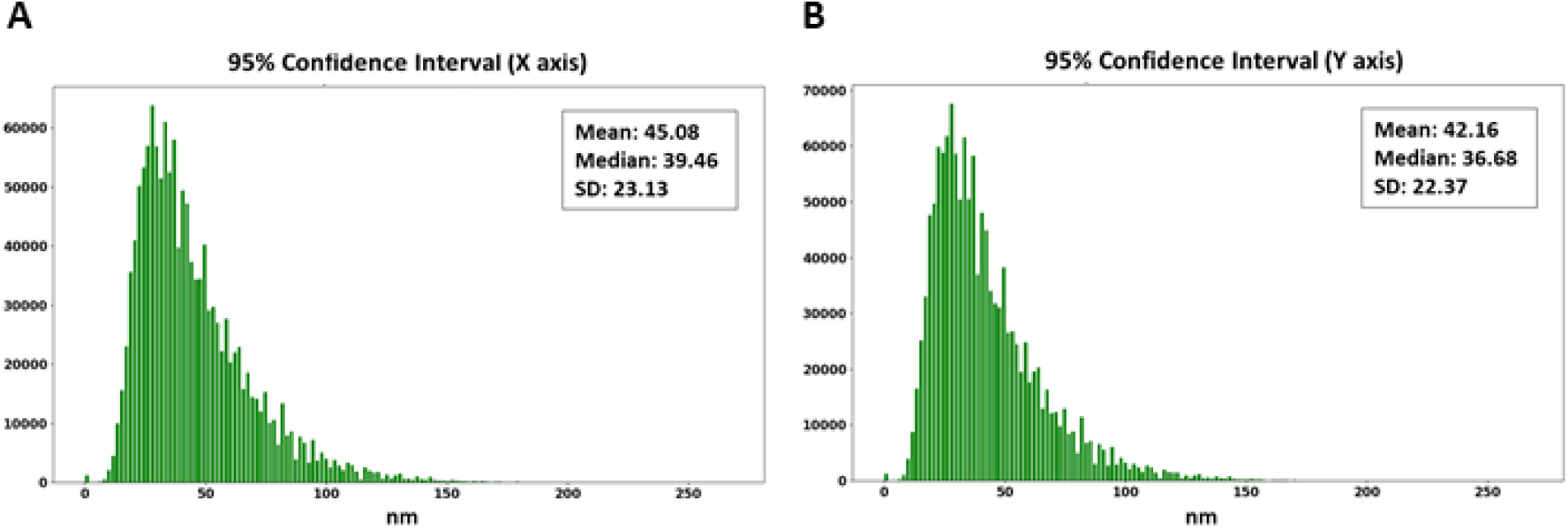
SMI Overall Localization Accuracy. The data shows an overall localization accuracy of 42 to 45 nm on average over the entire experiment in X and Y planes. The position of each transcript is sub-pixel localized using a 2D polynomial to estimate the X and Y coordinates of the transcript. The observed histogram of the uncertainty in X and Y coordinates of each called transcript over the course of an entire run is shown, along with the standard deviation (SD) of the histogram.

**Figure S3.**
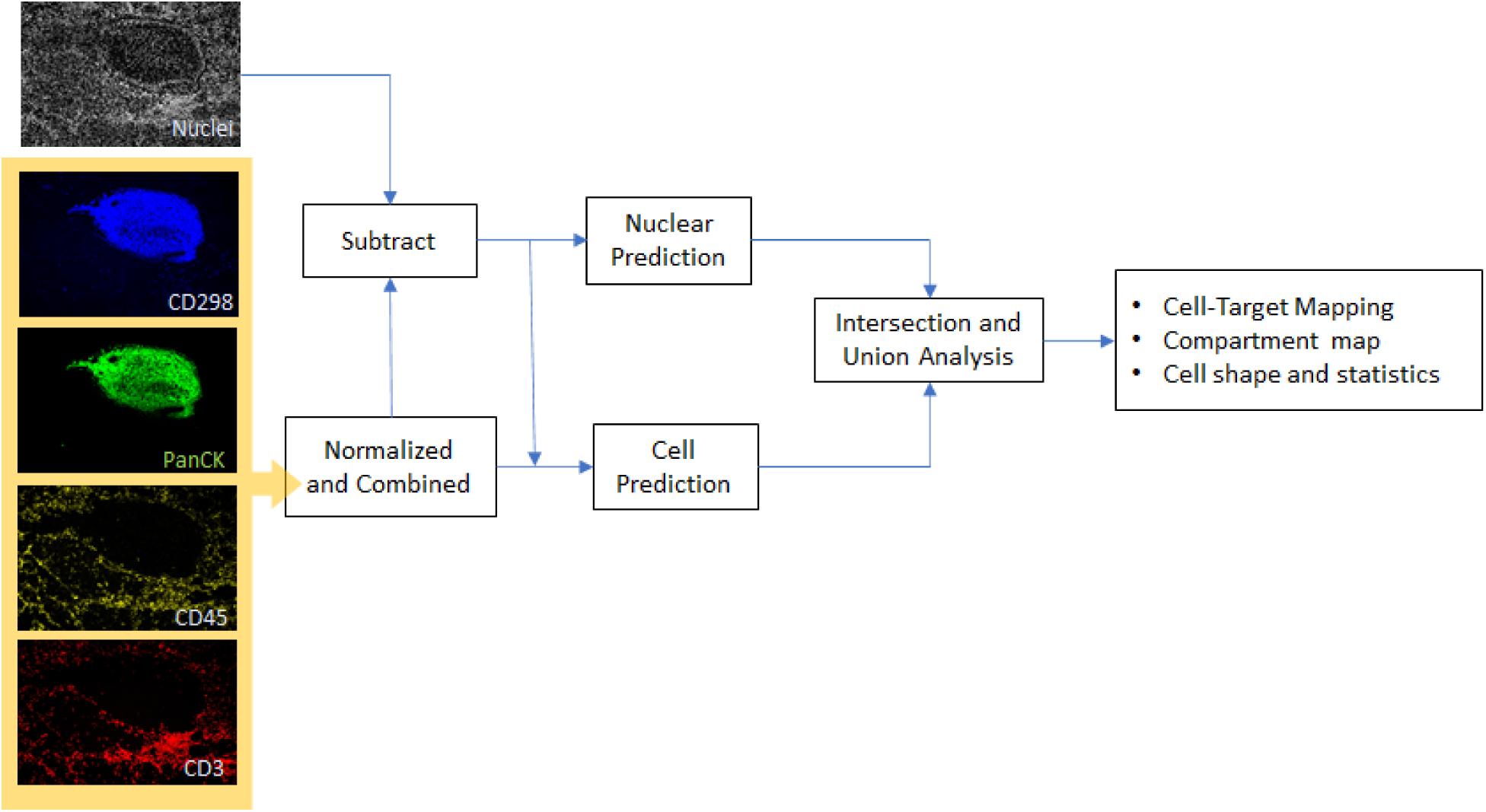
Cell Segmentation Pipeline. Membrane images are combined with normalization and subtracted with the nuclear image to enhance gaps between nuclei. Results are fed into the prediction network to segment the nuclei and cell boundaries. Results from segmentation are used to map the target to its corresponding cell and compartment within the cell.

**Figure S4.**
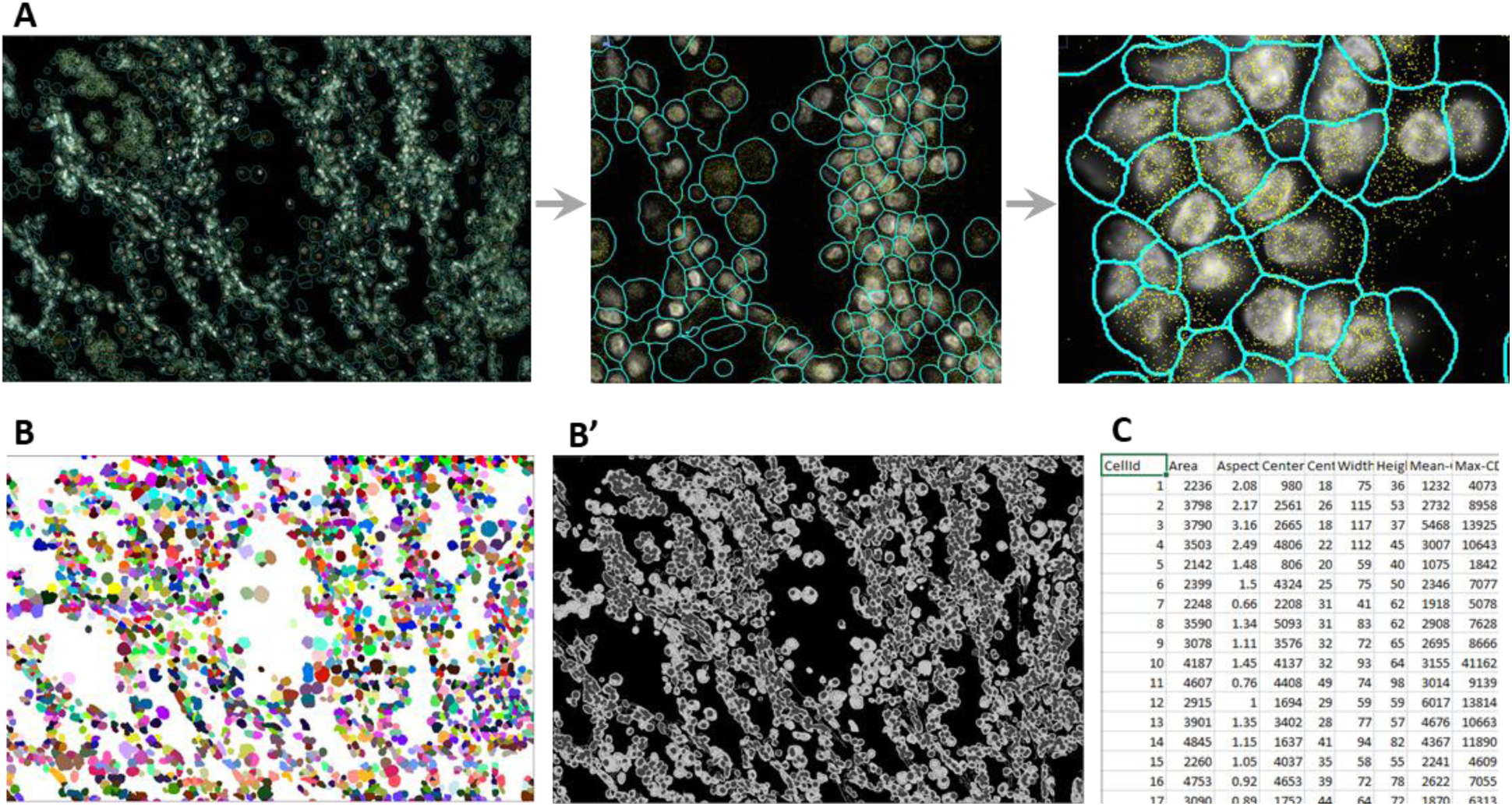
Cell Segmentation Results in Non-Small Cell Lung Carcinoma (NSCLC) Tissue. **A.** Segmentation overlay output: Segmentation boundaries (cyan) overlaid with nuclear image and transcripts (plotted in yellow) at different zoom levels. **B.** Cell label output: each cell is marked with unique cell ids (shown as different colors). **B’.** Cell compartment map: an output to show specific compartments of nuclei, membrane, and cytoplasm (marked with different gray level values). **C.** Cell statistics table output: area, aspect ratio, bounding box, and intensity statistics per channel are listed for each cell (id).

**Figure S5.**
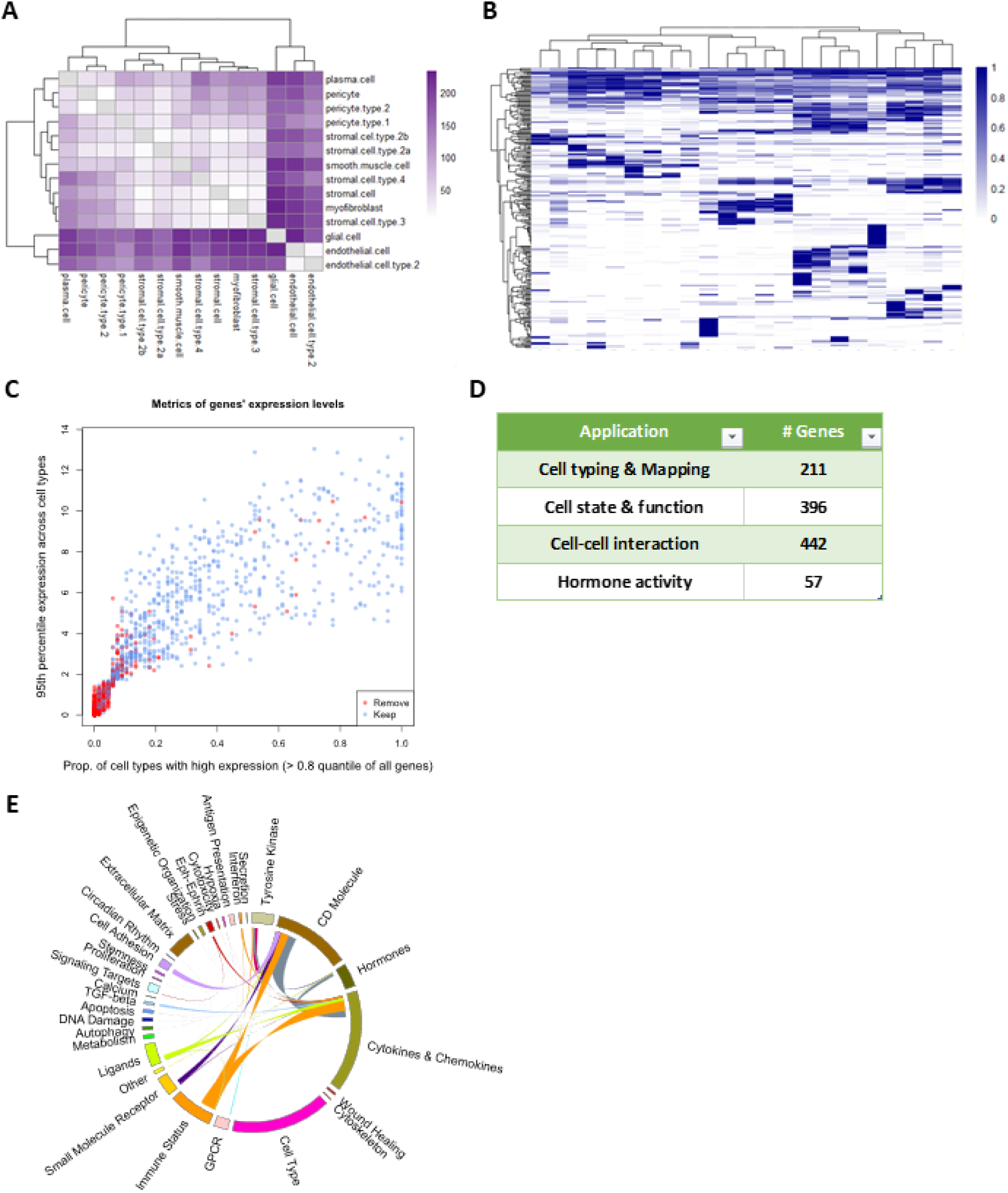
Content Design for a 980-plex Single Cell Panel for Measuring Cell Type, Cell State, and Cell-Cell Interactions. **A.** Example of starting point for gene selection, demonstrating which cell types were selected for additional discrimination after accounting for biologically relevant genes. **B.** Example of cell type profiles over complete gene list (per kidney scRNA-seq). Most genes discriminate between cell types (genes on y-axis, cell types on x-axis). **C.** Using scRNA-seq average cell type profiles from 76 Human Cell Landscape datasets, we scored genes across cell types based on the level of expression and consistency of expression across cell types. Genes selected based on scoring metrics are shown. Other genes were considered required based on strong biological relevance. **D.** Final gene counts by purpose; categories not mutually exclusive. **E.** Circos diagram summarizing the overlapping genes between the panel’s gene sets. Cell type is a set designed to increase the contrast between cell types beyond the biological content.

**Figure S6.**
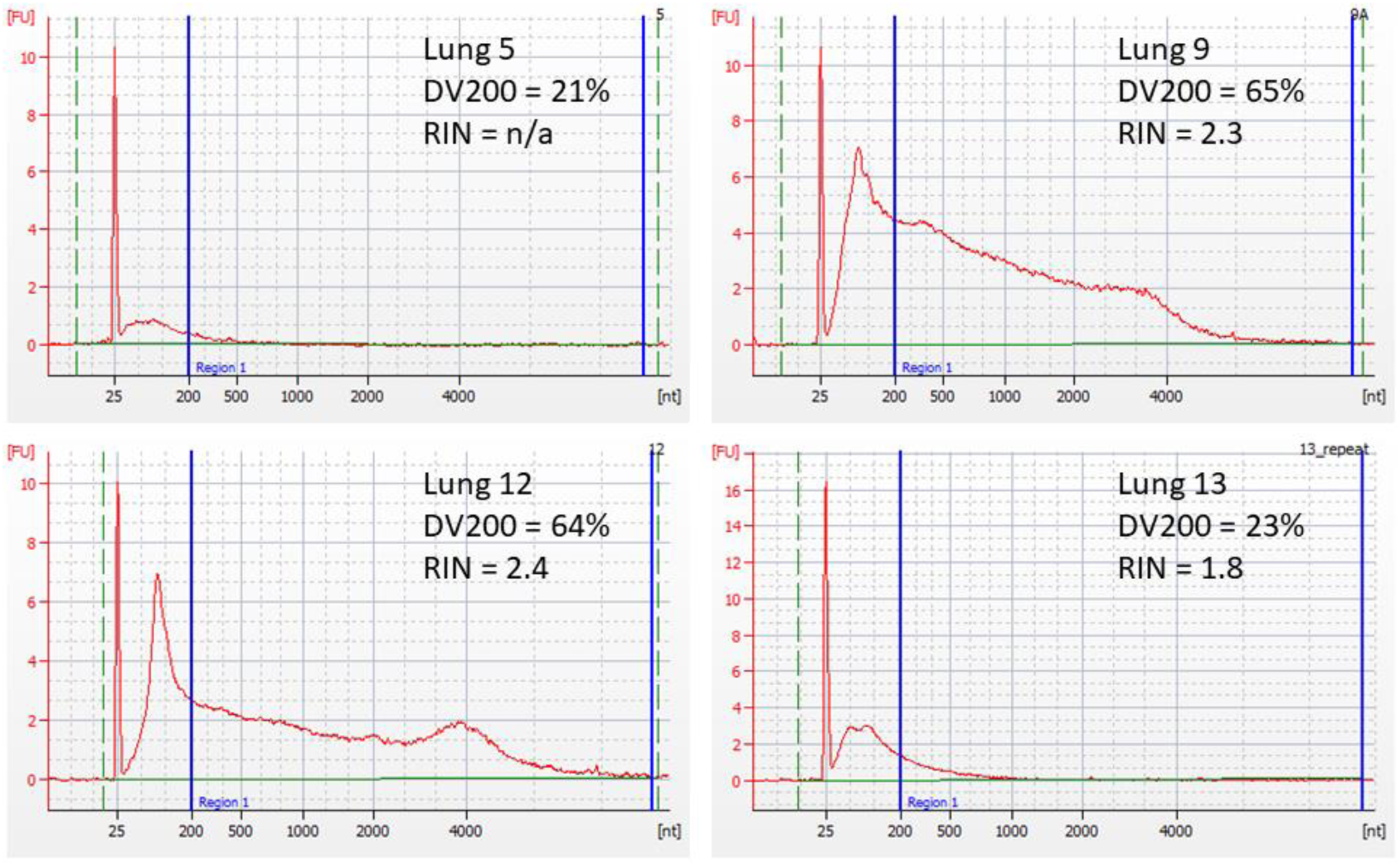
RNA Quality of Lung Samples. DV200 and RIN scores are determined using Agilent BioAnalyzer 2100 Expert Software B.02.09. For DV200, Region 1 selection is from 200 nt to about 8,000 nt using Smear Analysis.

**Figure S7.**
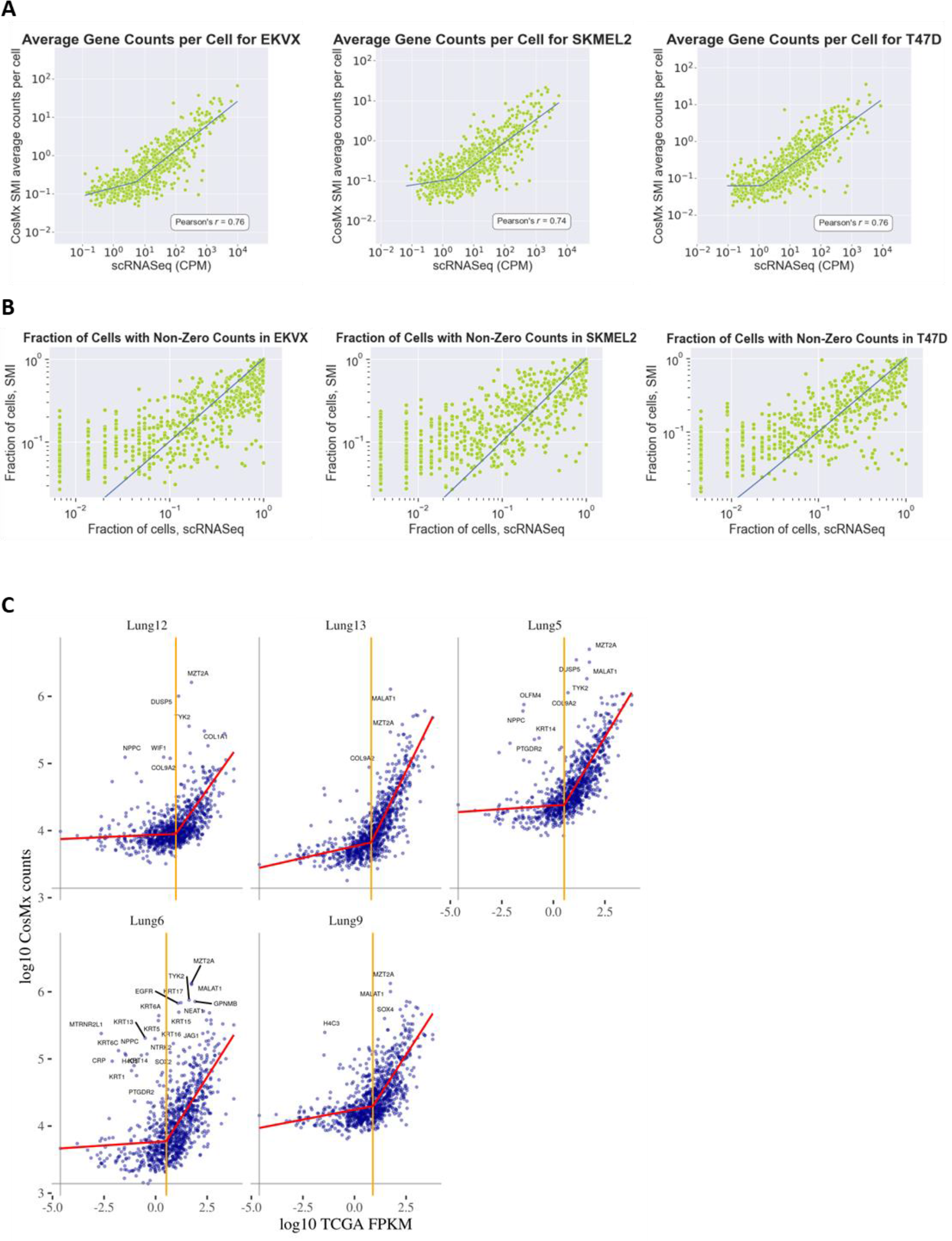
The correlation between SMI data and published scRNA-seq data from three matched cancer cell lines and bulk RNA-seq data from NSCLC. **A.** The counts per cell of SMI RNA assays are compared to published CPM-normalized scRNA-seq data (CPM) from lung adenocarcinoma (EKVX), melanoma (SKMEL2), and breast cancer (T47D) cell lines (38). **B.** Comparison of the number of cells per gene, calculated for the fraction of cells that have non-zero counts for each gene, between SMI and scRNA-seq data from these cell lines. **C.** The SMI bulk profile of each NSCLC sample is compared to the log_10_ mean FPKM of NSCLC in TCGA.

**Figure S8.**
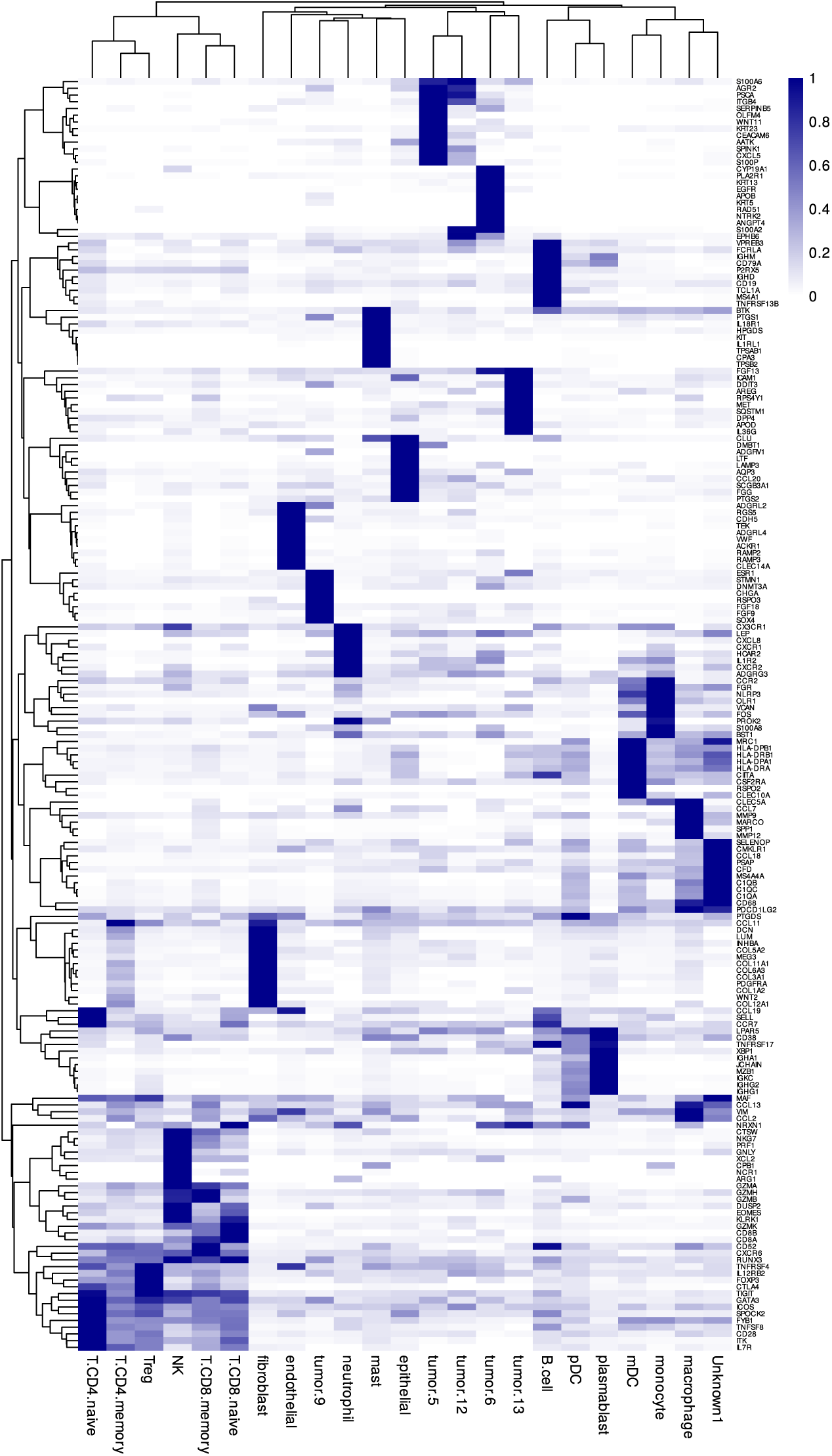
Reference Gene Expression Profiles Used in Supervised Cell Typing. Each row is scaled to have a maximum of 1.

**Figure S9.**
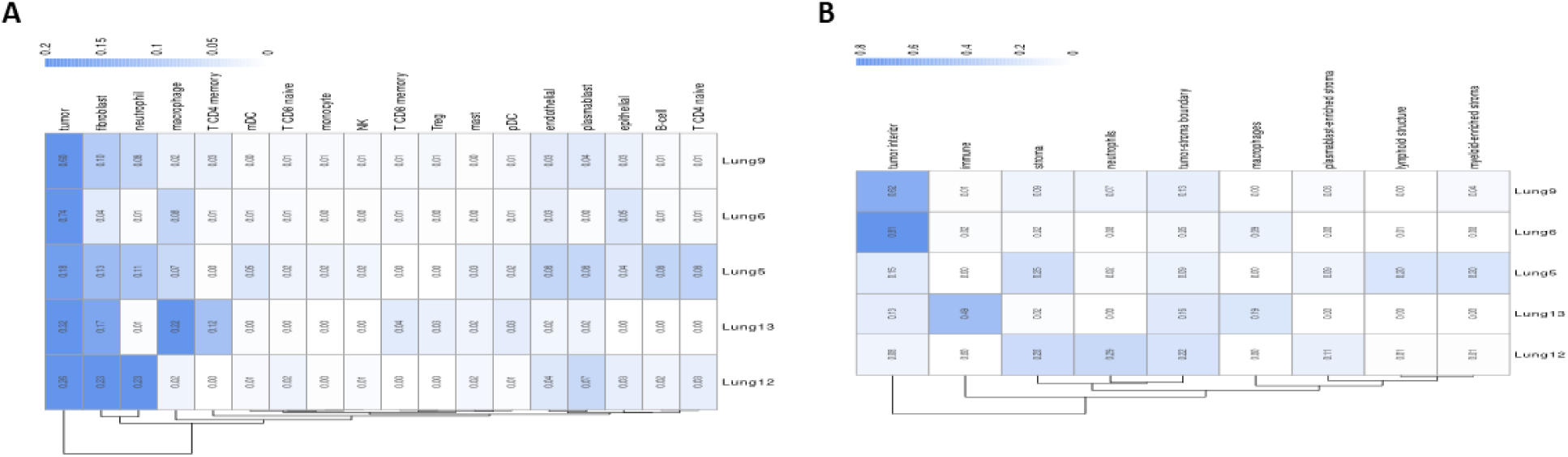
Extracting Tissue-Level Characteristics. **A.** Abundance of cell types within each tumor. **B.** Abundance of microenvironment niches within each tumor.

**Figure S10.**
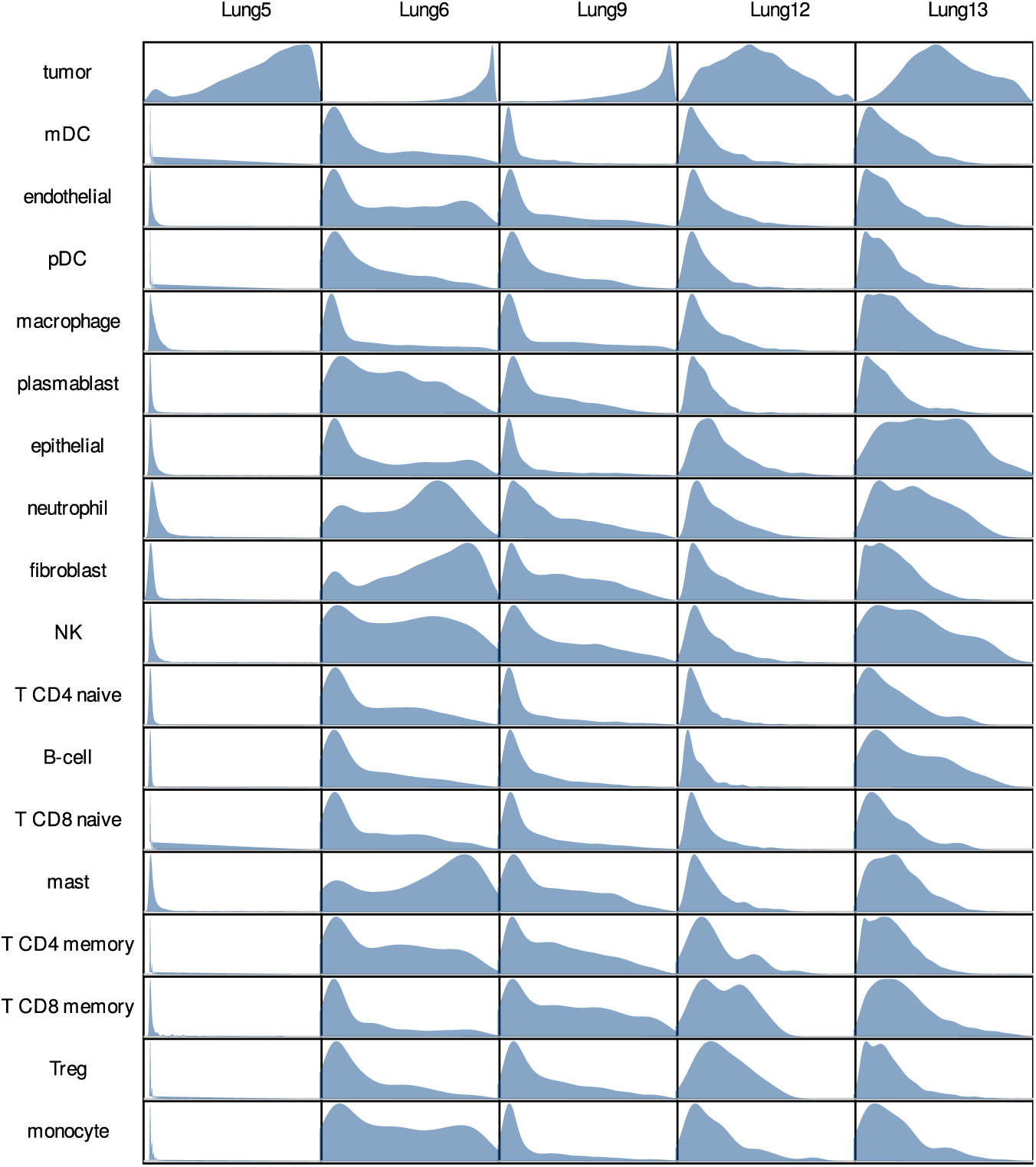
Invasiveness of Each Cell Type Within Each Tissue. Cells are scored for the frequency of tumor cells within their 100 nearest neighbors. Shapes show the density of the score within each cell type.

**Figure S11.**
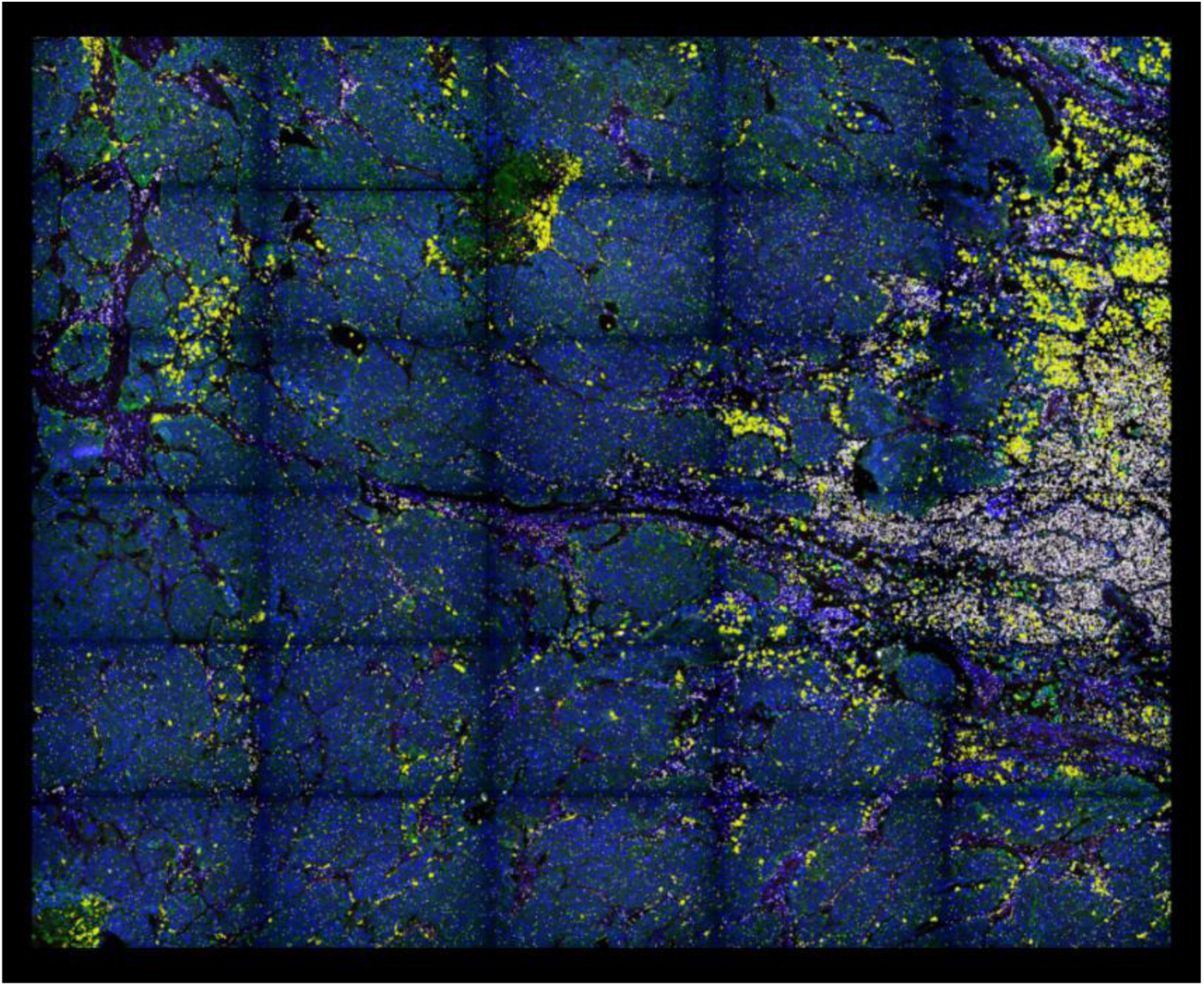
Highlighting Transcripts of SPP1 and HLA-DQA1 in Lung 6. SPP1 transcripts are shown as yellow dots; HLA-DQA1 transcripts are shown as white dots.

**Figure S12.**
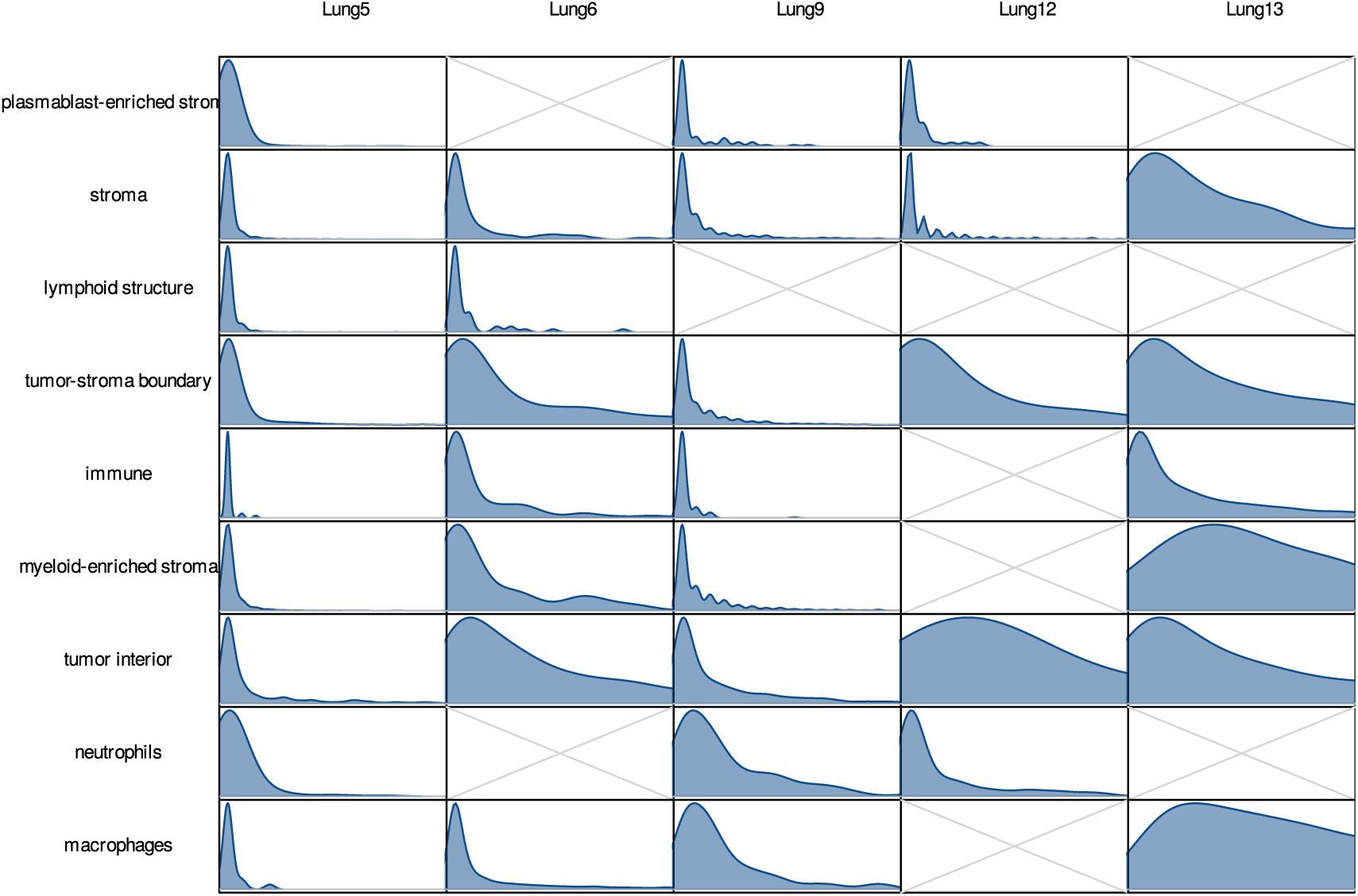
Expression of SPP1 Across Tumors and Niches. Polygons show densities of SPP1 transcripts per macrophage; horizontal axes are truncated at 10 counts. Only tumors and niches with >20 macrophages are shown.

**Figure S13.**
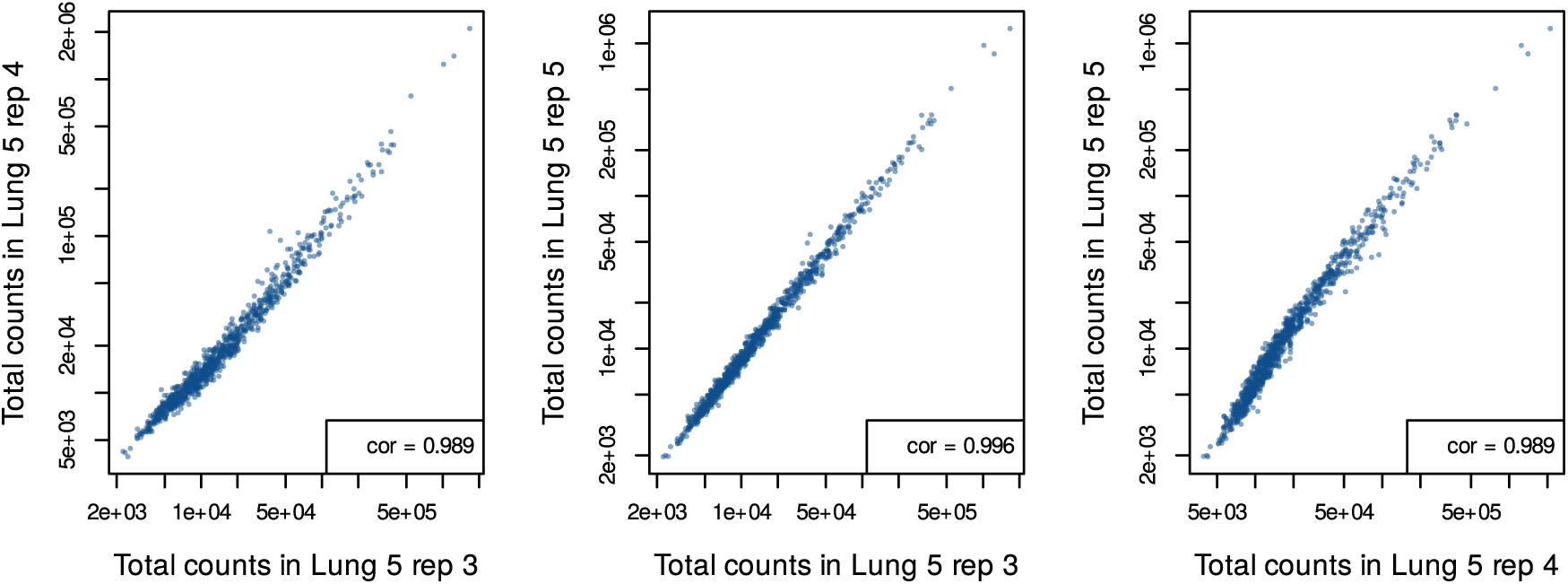
Concordance of Bulk Expression Profiles from Three Replicates. For each of the three replicates, total transcripts of each gene were recorded, and these bulk profiles are compared across replicates. Correlation is calculated on log-transformed data.

**Figure S14.**
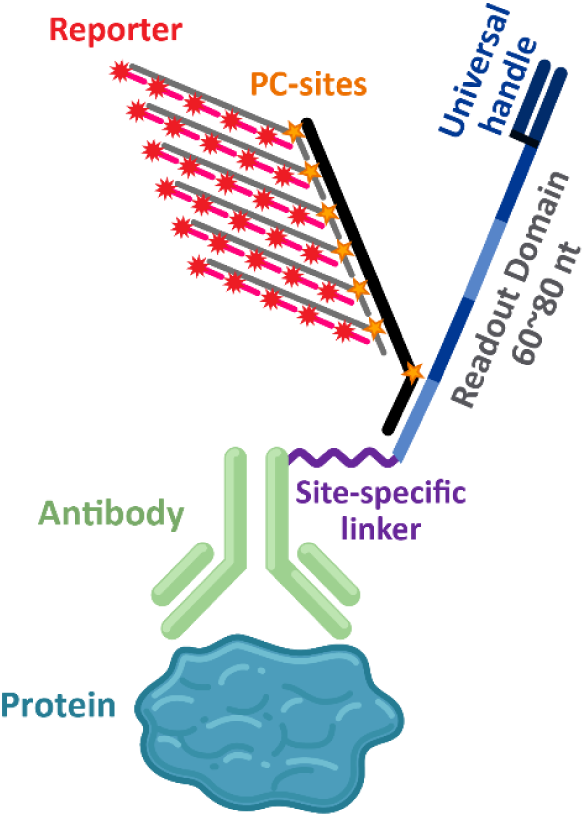
Protein Detection Probe. The protein detection probe is readily adapted from the RNA detection probe shown in Figure 1. For protein detection, the readout domain contains an additional universal handle and is conjugated to the antibody via a site-specific linker.

**Figure S15.**
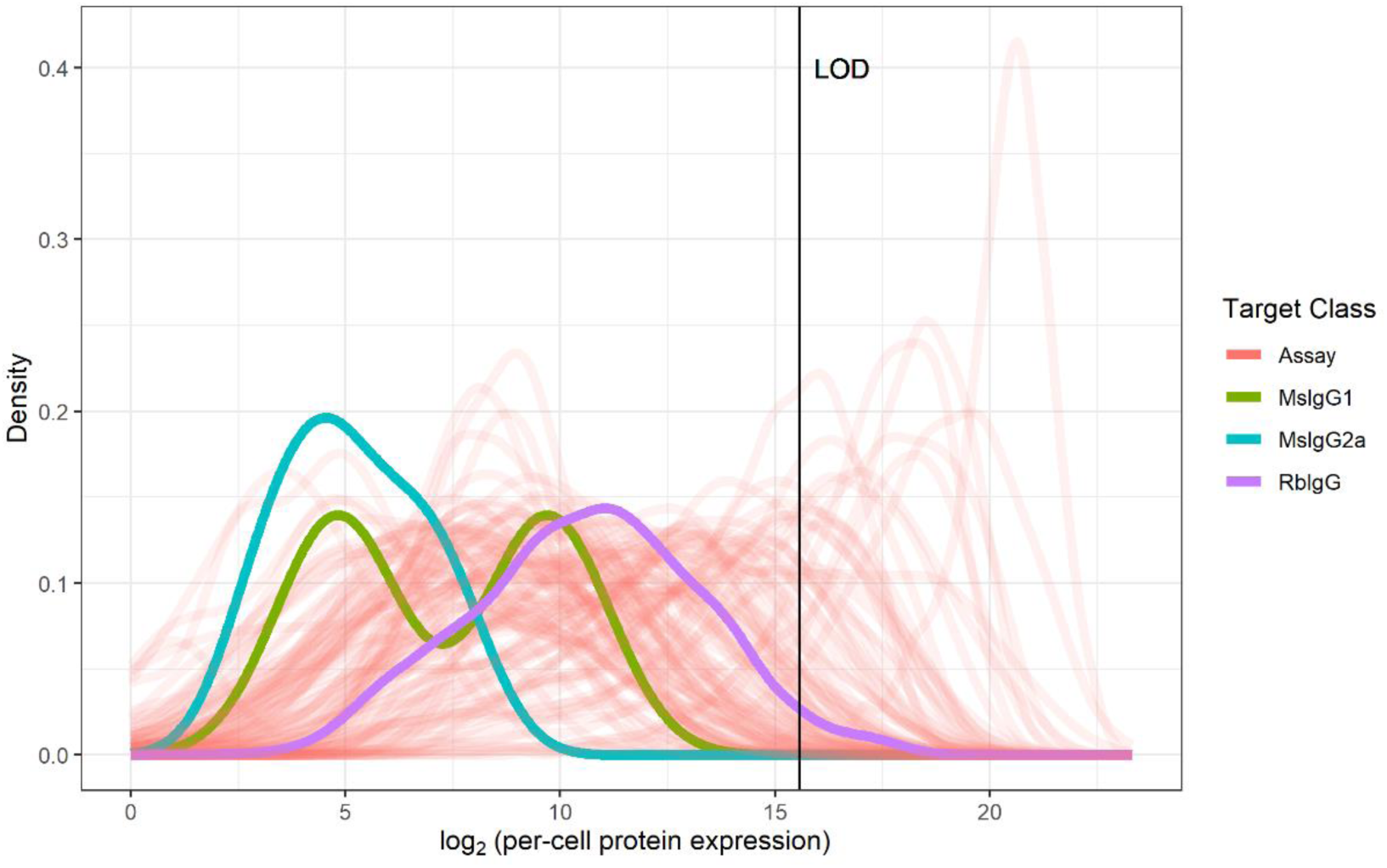
Distribution of Per-Cell Expression Values for Targets in the Protein Assay. The distribution of per-cell protein expression values for the FOV visualized in Figure 6 is colored by the encoded targets in the assay (“Assay”) or one of three negative isotype controls. The limit of detection (LOD), set for visualization of single cell expression values, is plotted as a black vertical line. The LOD was set by filtering to cells with detectable signal from at least one negative control, taking the geometric mean of per-cell expression and adding 1.5 standard deviations. As indicated by the LOD position relative to the density profile for one of three isotype controls, and as evident in Figure 6C, 51 of 2688 cells in this FOV had rabbit IgG levels above the LOD. MsIgG1: mouse IgG1, MsIgG2a: mouse IgG2a, and RbIgG: rabbit IgG.

**Figure S16.**
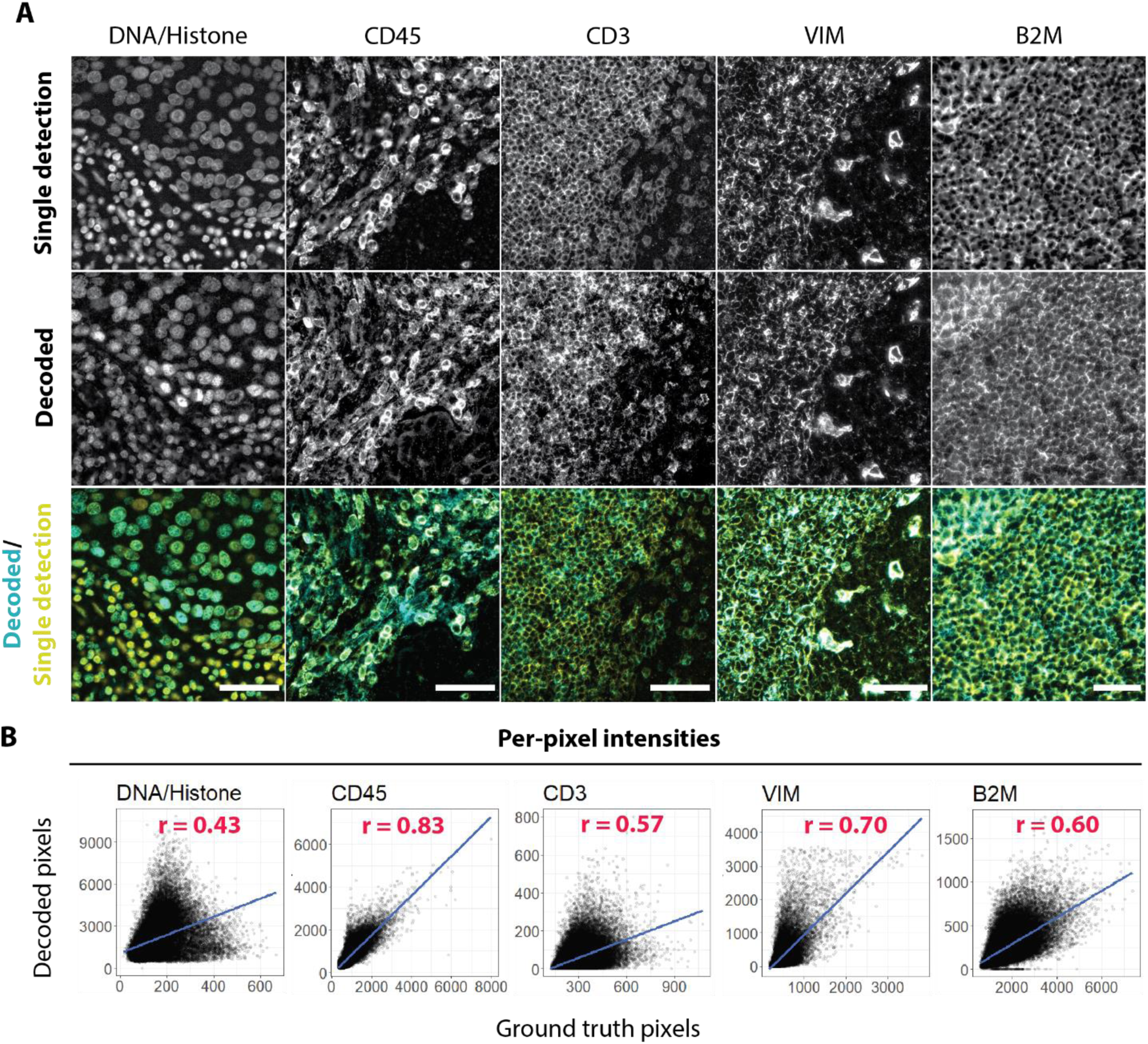
Concordance Between Protein Localization Patterns Detected in Multiplexed SMI Protein Assay with Ground Truth Data. **A**. Visualization of protein localization obtained from traditional single-channel immunofluorescence detection compared to that obtained from decoding by the SMI protein assay. Single detection of DNA was performed with DAPI staining. DNA/Histone and CD45 experiments were performed on lung cancer tissue. CD3, vimentin (VIM), and B2M experiments were performed on tonsil tissue. Ground-truth CD45 and B2M were imaged using a morphology marker and reporter pair. Ground-truth CD3 and VIM were obtained using fluorescently conjugated antibodies following the completion of the SMI readout with on-instrument staining. Ground truth and decoded data for CD45 and B2M were obtained from antibodies of the same clone. Data for CD3 and VIM were obtained from antibodies that recognized different epitopes. Scale bar: 50 µm. **B**. Per-pixel spatial correlation between ground truth pixels (from single-detection image) and decoded pixels (from decoded image). 50,000 pixels were randomly subsampled from 1 million pixels and plotted against their spatially corresponding pixels. A standard linear regression line is shown in blue. Pearson R correlation coefficient is reported for each protein target (red).

**Figure S17.**
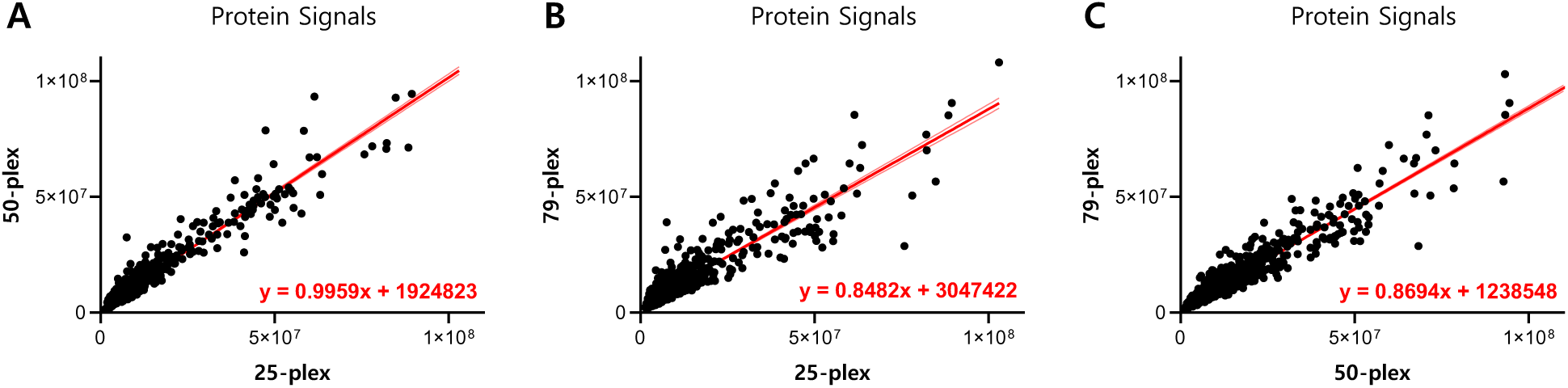
SMI Protein Assay Delivers Consistent Results as the Plex Size Increases. Relative protein abundance in 35 different cell lines was evaluated by the SMI protein assay. The number of antibodies simultaneously detected ranged from 25 to 79 antibodies. Our algorithm used the same encoding scheme to decode all 3 datasets. Protein expression values for each cell line were normalized by cell count, and pairwise comparisons of shared targets were conducted between: **A.** 25-plex and 50-plex assays, **B.** 25-plex and 79-plex assays, and **C.** 50-plex and 79-plex assays. Total pixel intensity is used as a relative unit of measurement for protein abundance. A standard linear regression line is shown in red. Pearson R correlation coefficient and linear regression fit function are reported for each pairwise comparisons (red).

**Figure S18.**
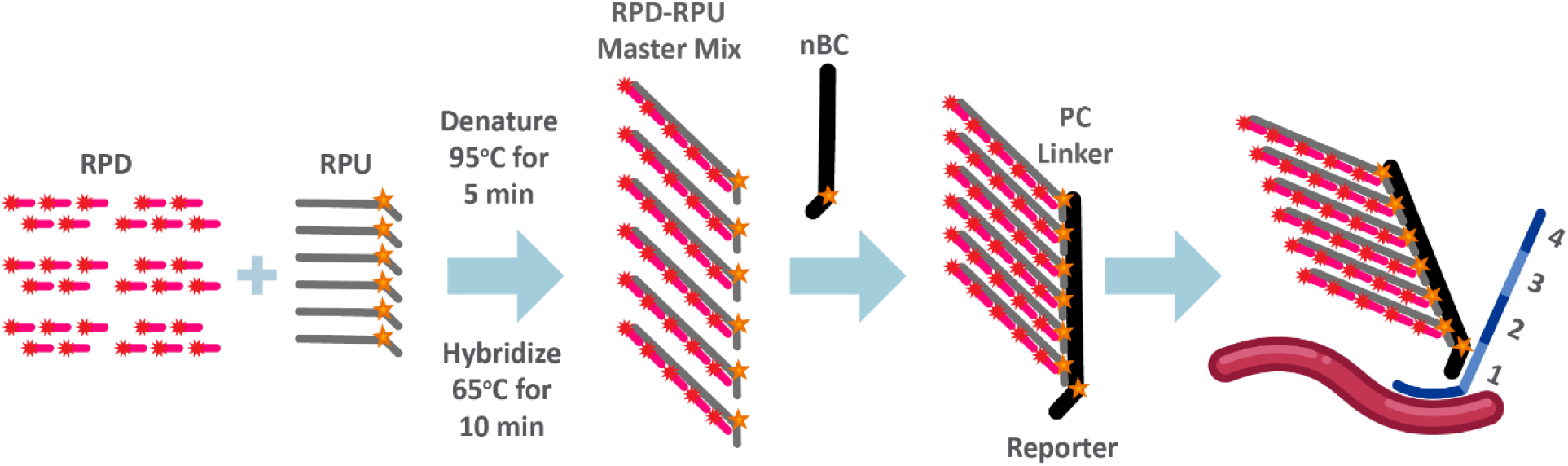
Reporter Readout Probe Design and Assembly. Reporter probe dye (RPD) and sub-branch (RPU) are denatured to ensure all molecules are unhybridized and then allowed to hybridize to produce the RPD-RPU Master Mix. This Master Mix is hybridized to the nanoBarCode (nBC) to form a single-color reporter. These reporters are pooled by the hybridization region in the probe. Each set of pools contains four reporters with four different fluorophores (*e.g*., blue, red, yellow, green reporters for spot 1).

